# Norovirus NS1/2 protein increases glutaminolysis for efficient viral replication

**DOI:** 10.1101/2023.12.19.572316

**Authors:** Adam Hafner, Noah Meurs, Ari Garner, Elaine Azar, Karla D. Passalacqua, Deepak Nagrath, Christiane E. Wobus

## Abstract

Viruses are obligate intracellular parasites that rely on host cell metabolism for successful replication. Thus, viruses rewire host cell pathways involved in central carbon metabolism to increase the availability of building blocks for replication. However, the underlying mechanisms of virus-induced alterations to host metabolism are largely unknown. Noroviruses (NoVs) are highly prevalent pathogens that cause sporadic and epidemic viral gastroenteritis. In the present study, we uncovered several strain-specific and shared host cell metabolic requirements of three murine norovirus (MNV) strains, the acute MNV-1 strain and the persistent CR3 and CR6 strains. While all three strains required glycolysis, glutaminolysis, and the pentose phosphate pathway for optimal infection of macrophages, only MNV-1 relied on host oxidative phosphorylation. Furthermore, the first metabolic flux analysis of NoV-infected cells revealed that both glycolysis and glutaminolysis are upregulated during MNV-1 infection of macrophages. Glutamine deprivation affected the MNV lifecycle at the stage of genome replication, resulting in decreased non-structural and structural protein synthesis, viral assembly, and egress. Mechanistic studies further showed that MNV infection and overexpression of the MNV non-structural protein NS1/2 increased the enzymatic activity of the rate-limiting enzyme glutaminase. In conclusion, the inaugural investigation of NoV-induced alterations to host glutaminolysis identified the first viral regulator of glutaminolysis for RNA viruses, which increases our fundamental understanding of virus-induced metabolic alterations.

**Author Summary:** All viruses critically depend on the host cells they infect to provide the necessary machinery and building blocks for successful replication. Thus, viruses often alter host metabolic pathways to increase the availability of key metabolites they require. Human noroviruses (HNoVs) are a major cause of acute non-bacterial gastroenteritis, leading to significant morbidity and economic burdens. To date, no vaccines or antivirals are available against NoVs, which demonstrates a need to better understand NoV biology, including the role host metabolism plays during infection. Using the murine norovirus (MNV) model, we show that host cell glutaminolysis is upregulated and required for optimal virus infection of macrophages. Additional data point to a model whereby the viral non-structural protein NS1/2 upregulates the enzymatic activity of glutaminase, the rate-limiting enzyme in glutaminolysis. Insights gained through investigating the role host metabolism plays in MNV replication may assist with improving HNoV cultivation methods and development of novel therapies.

## Introduction

Viruses are metabolically inert and must rely on host cell metabolic events to generate the necessary building blocks to multiply (1). Historically, host metabolism has been thought to play only host-specific roles in cellular homeostasis, the immune response, and autophagy (2–3). However, recent studies have shown that pathogens such as parasites, bacteria, and viruses influence host cell metabolism (4–6) to create a more favorable environment to ensure their own optimal replication (15). Many investigations within the past decade have examined how viruses alter the host cellular metabolic profile and identified some of the metabolic pathways important during virus infection. These studies have shown that a common consequence of viral infection is induction of high glucose metabolism, which can lead to aerobic glycolysis, or the Warburg effect (7). In addition, other pathways such as glutaminolysis, the pentose phosphate pathway (PPP), fatty acid synthesis, and tricarboxylic acid cycle (TCA) activity may also be altered, thus highlighting that central carbon metabolism is significantly perturbed during many viral infections (7). Viruses often hijack these pathways to divert the production of nucleotides, lipids, amino acids, and other metabolites away from host processes toward virus particle construction. Virus-induced alterations to host metabolism can be shared among different viruses but are usually context dependent and variable between specific virus families or infected host cell types. For example, glucose deprivation significantly decreases dengue virus replication, while lack of glutamine does not (8). In contrast, glutamine deprivation significantly reduces vaccinia virus replication, while glucose deprivation has no effect (9). Thus, dengue virus and vaccinia virus show opposite dependencies on host glycolysis and glutaminolysis during infection. Other examples of virus-induced changes in host metabolism come from adenovirus, human cytomegalovirus, chikungunya virus, Zika virus, SARS-CoV-2, rhinovirus, lytic gammaherpesvirus, both latent and lytic Kaposi sarcoma-associated herpes virus and hepatitis C virus (10–14, 33, 36, 38–39, 68). While multiple studies have reported that metabolic pathways are altered during virus infection, the mechanistic details of how viruses achieve these changes remain elusive. Increased investigation into how viruses reprogram and usurp host metabolic pathways with an emphasis on mechanistic insights may reveal innovative therapeutic targets and provide a deeper understanding of specific viral replicative cycles.

Noroviruses (NoVs) are positive-sense single-stranded RNA viruses and the leading cause of acute non-bacterial gastroenteritis worldwide (16). Globally, human NoV (HNoV) infections are extremely common, with estimated cases reaching ∼685 million per year. Annually, HNoV infections result in ∼200,000 fatalities, mostly in infants but also in immunocompromised individuals and in older adults (17). Additionally, HNoV infections result in serious annual economic burdens, with global economic costs surpassing US$60 billion (18). In the United States alone, HNoV infections cause ∼21 million cases of gastroenteritis and are the leading cause of death in older adults with viral gastroenteritis (19–20). Although HNoV infections are self-limiting in most individuals, the intense vomiting, diarrhea, and abdominal pain associated with this infection can be debilitating. However, despite the devastating public health and economic burdens caused by HNoV, no approved vaccines or antivirals against this virus exist (21), and development of anti-NoV therapeutics has been hampered by the lack of a cell culture model for HNoV. Although human intestinal enteroids (HIEs) and human B cells support varying degrees of infection, a cell culture–derived HNoV stock is still not available (22–24). To overcome the limitations inherent to HNoV research, murine NoV (MNV) is used as a model system to study general NoV biology because MNV readily replicates in cell culture, is genetically similar to HNoV, and has a genetically tractable small animal model and infectious clones available (25). MNV strains, although genetically closely related, fall into two phenotypic groups. The acute strain, MNV-1, is cleared from infected mice within one week, while persistent strains, including MNV-CR6 (CR6) and MNV-CR3 (CR3), are shed for months (26). The strains also differ in their *in vivo* tropism, in which CR6 infects tuft cells while MNV-1 infects immune cells (macrophages, dendritic cells, and lymphocytes) (27,28).

We previously performed a metabolomic screen of MNV-1–infected macrophages, which revealed that metabolites in many pathways were significantly upregulated, including those integral to central carbon metabolism (29). Our screen identified glycolysis, nucleotide biosynthesis via the PPP, and oxidative phosphorylation (OXPHOS) as being required for optimal MNV-1 replication in murine macrophages based on experiments using common metabolic inhibitors (29). We further determined that glycolysis is important for the replication step in the MNV lifecycle since treatment with the hexokinase inhibitor 2-deoxyglucose (2DG) led to a decrease in viral protein and RNA synthesis (29). However, the requirement for glycolysis was independent of the host antiviral type I interferon response, and the underlying mechanisms behind NoV-induced upregulation of host metabolism and the role that host metabolic pathways plays in persistent MNV replication are not known. Thus, the goals of this current study were to further define the role of host metabolism in NoV replication, explore the role of host metabolism for persistent MNV strains, and begin to uncover the underlying mechanisms of NoV-induced metabolic alterations. Untangling the process of virus-induced metabolic alterations may enable development of more efficient HNoV cultivation systems and identify innovative metabolic therapeutic targets aimed at reducing persistent NoV infections.

With these goals in mind, we investigated the dependence of persistent strains CR3 and CR6 on host cell glycolysis, the PPP, and OXPHOS. While MNV-1, CR3, and CR6 all relied on glycolysis and nucleotide biosynthesis, OXHPOS was not required for replication of persistent strains. We also performed the first metabolic flux analysis of MNV-1–infected macrophages, which revealed a concurrent increase in glycolysis and glutaminolysis. Reducing host glutaminolysis via pharmacological inhibition with the inhibitor CB839 and via glutamine deprivation showed that both acute and persistent MNV strains rely on glutamine metabolism, in particular for viral genome replication, which has repercussions for later steps in the viral life cycle. Early mechanistic investigations revealed that the observed increase in glutaminolysis during MNV infection is driven in large part by the viral non-structural protein NS1/2 that caused increased glutaminase (GLS) activity, the rate limiting enzyme within the glutamine catabolic pathway (30). Overall, our findings highlight the importance of pathways in central carbon metabolism in NoV infection, albeit with strain-specific differences, and show that glutaminolysis is universally required for optimal MNV replication. Our finding that glutaminolysis is modulated by the viral protein NS1/2 provides a foundation for detailed mechanistic studies in the future, which may reveal novel chokepoints for therapeutic intervention.

## Results

### Persistent MNV strains CR6 and CR3 rely on glycolysis and nucleotide biosynthesis, but not OXPHOS, for optimal replication

We previously performed a metabolomics screen of MNV-1–infected macrophages, which identified increased metabolites from glycolysis, PPP, and OXPHOS in infected cells (29). Inhibition of these pathways resulted in significantly lower MNV titers, ranging from an 0.5 to 2- log_10_ reduction (29). However, whether the genetically closely related persistent MNV strains CR3 and CR6 also rely on these important metabolic pathways for optimal replication was not known. To investigate whether acute and persistent MNV strains have a common dependence on host cell metabolism, RAW 264.7 (RAW) cells were inoculated with MNV-1, CR3, and CR6 at an MOI of 5 for 1 hour. Medium containing the glycolysis inhibitor 2DG, the PPP inhibitor 6-Aminonicotinamide (6AN), or the OXPHOS inhibitor oligomycin-A was then added after inoculation, and cells were incubated for 8 hours, corresponding to approximately one round of viral replication. Non-toxic concentrations of 2DG and 6AN were previously determined (29), and cell viability assays were performed to ensure the concentration of oligomycin-A used would maintain >80% cell viability (Fig S1A). Infectious titers were measured after 8 hours via plaque assay. A significant (>2 log_10_) decrease was observed in the number of infectious MNV-1, CR3, and CR6 titers in 2DG-treated cells (Fig. 1A). Treatment with 6AN also resulted in significantly decreased MNV-1, CR3, and CR6 titers; however, only a 1 log_10_ decrease in infectious particles was observed (Fig 1B). Additionally, the 1 versus 2 log_10_ decrease in viral titers observed after 6AN and 2DG treatment, respectively, suggested that all three MNV strains depend more on glycolysis than the PPP for optimal reproduction.

**Figure 1:**
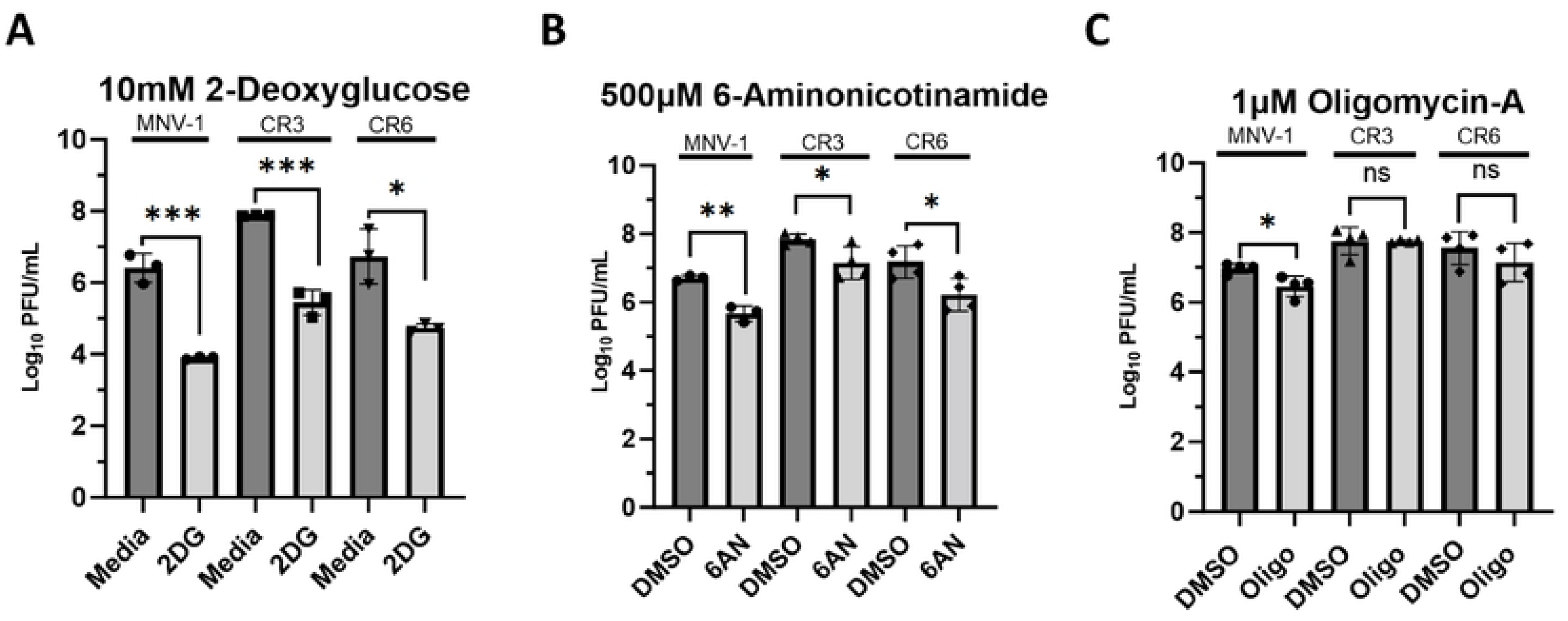
Persistent strains CR3 and CR6 rely on host glycolysis and nucleotide biosynthesis, but not OXPHOS, for optimal replication. RAW 264.7 cells were infected for 1 hour at an MOI of 5 with either MNV-1, CR3, or CR6. Virus inoculum was removed and replaced with medium containing **(A)** 10 mM 2-deoxuglucose (2DG), **(B)** 500 μM 6-aminonicotinamide (6AN), **(C)** 1 μM oligomycin-A (Oligo), or vehicle control (DMSO). Infected cells were incubated for 8 hours and infectious MNV titers were measured via plaque assay. Experiments represent combined data from at least three independent experiments. Statistical analysis was performed using Two-tailed Students-t tests. ***, *P<*0.001; **, *P* <0.01; *, *P<*0.05; ns, not significant.

Because active viral replication requires large amounts of host energy, we also investigated whether CR3 and CR6 require OXPHOS for optimal replication. Surprisingly, we observed that CR3 and CR6 infection did not depend on OXPHOS because viral titers remained similar between oligomycin-A treated and untreated cells; however, acute strain MNV-1 showed an 0.5-log_10_ titer decrease. Lack of a significant reduction in viral titers of persistent MNV strains during oligomycin-A treatment suggests that glycolysis-derived ATP is sufficient to meet the energetic requirements for sustaining optimal CR3 and CR6 replication. These data highlight strain-specific dependencies on individual metabolic pathways for efficient MNV virion production.

Taken together, these data demonstrate that like MNV-1, the persistent strains CR3 and CR6 require host glycolysis and nucleotide biosynthesis for optimal replication; however, unlike MNV-1, OXPHOS is dispensable for the persistent strains.

### MNV-1 infection upregulates metabolite flux through glycolysis and glutaminolysis

Our previous static metabolomic screen (29) analyzed the intracellular concentrations of metabolites but did not measure metabolite flux or metabolite turnover. To this end, we performed a metabolic flux analysis, which uses uniformly labeled metabolites measured via gas chromatography mass spectrometry (GC-MS) to track incorporation of molecules into various metabolic pathways. Because glucose and glutamine are the two leading carbon sources used by mammalian cells (31), we analyzed their incorporation during MNV-1 infection to determine whether infection mediates an increase in their catabolism (Fig. 2). RAW cells were infected for 1 hour with MNV-1 or mock lysate at an MOI of 5. After a 1-hour incubation, the virus inoculum was replaced with medium containing either ^13^C_5_-glucose or ^13^C_5_-glutamine. Samples were collected and analyzed after an 8-hour incubation. Through analysis of the mass isotopomer distribution (MID), we observed higher glucose metabolism in MNV-1–infected cells than in mock-infected cells as seen by increased incorporation of glucose into lactate, a common glycolytic byproduct, and into citrate, a downstream metabolite within the TCA cycle that can be generated from the final glycolytic product pyruvate through acetyl co-enzyme A (Fig. 2A). These findings are consistent with our previous metabolic screen that showed higher concentrations of several glycolytic intermediates such as 2- and 3- phosphoglycerate and fructose-bisphosphate in infected cells (29) and confirmed that MNV-1 induces host glucose metabolism during its replicative cycle. Additionally, we further observed increased glutamine metabolism in MNV-1–infected cells relative to mock-infected cells. Glutamine undergoes a deaminase reaction to produce glutamate followed by another deaminase reaction to produce alpha-ketoglutarate (aKG), an intermediate that can enter the TCA cycle (Fig. 2B). In MNV-1–infected cells, higher production of both metabolites was observed, thus showing increased glutamine metabolism (Fig. 2B). Given this finding, we revisited our previous metabolomic screen and investigated whether the concentrations of glutamate or aKG were significantly altered during MNV-1 infection. While aKG was not included in the screen, glutamate levels were significantly higher during infection (29). Taken together, our previous metabolomic screen (29) and current flux analysis provide strong evidence that glutamine metabolism is upregulated during MNV infection. As a control to ensure that the presence of uniformly labeled glucose and glutamine did not negatively affect virus replication, we titered MNV-infected RAW cells in the presence of the labeled metabolites and measured viral replication via plaque assay (Fig. 2C). We observed no negative effects from the uniformly labeled metabolites on virus replication, with a >6 log_10_ growth after 8 hours (Fig. 2C), which is similar to titers obtained in unlabeled medium (Fig. 1).

**Figure 2:**
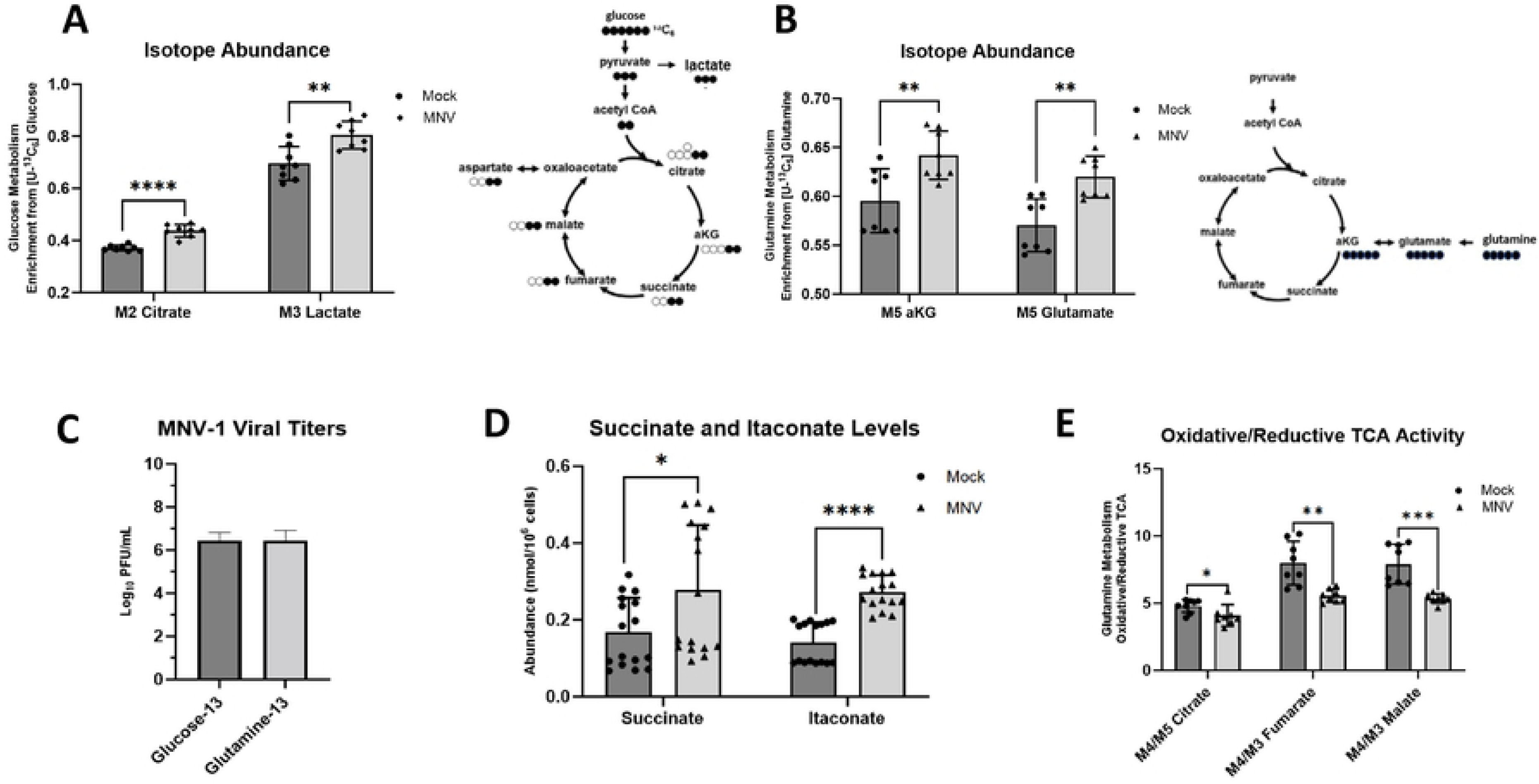
MNV-1 infection upregulates glycolysis and glutaminolysis in macrophages. RAW 264.7 cells were mock-infected or infected with MNV-1 for 1 hour at an MOI of 5. The virus inoculum was removed and replaced with medium containing (**A**) ^13^C_5_-glucose or (**B-E**) ^13^C_5_-glutamine for 8 hours. After 8 hrs, intracellular metabolites were extracted with ice-cold methanol. (**D**) RAW 264.7 cells were infected as before, and MNV-1 titers measured via plaque assay. Experiments represent combined data from at least two independent experiments with at least two technical replicates. Statistical analysis was performed by multiple unpaired t-tests. ****, *P<*0.0001; ***, *P<*0.001; **, *P* <0.01; *, *P<*0.05; ns, not significant.

Activated macrophages can dramatically upregulate immunoresponsive gene 1 (IRG1) expression leading to itaconate production from cis-aconitate in the TCA cycle (74–76). Furthermore, itaconate can play diverse roles in the immune response, including inhibition of succinate dehydrogenase in the TCA cycle (77). Consistent with previous reports, we measured approximately two-fold higher itaconate and succinate abundances (Fig. 2D) with a larger fraction being glutamine-derived in MNV-1 infected cells (Fig. S2A). To determine how itaconate production might affect mitochondrial metabolism in macrophages, we analyzed the utilization of reductive carboxylation in MNV-1 infected cells. Reductive carboxylation is a glutamine-dependent metabolism favored by cells when the oxidative mitochondrial metabolism is dysfunctional (69). We reasoned that production of itaconate during viral infection may reduce reliance on oxidative metabolism. Indeed, we measured a decrease in the ratio of oxidative to reductive metabolism in MNV-1 infected cells as measured by the ratio of oxidative-derived M4 citrate, M4 fumarate, and M4 malate to reductive-derived M5 citrate, M3 fumarate, and M3 malate (Fig. 2E).

Overall, flux analysis of MNV-1 infection demonstrates production of itaconate coupled with reductive TCA cycle activity and reprogramming of glucose and glutamine metabolism, which are all hallmarks of virus-induced metabolic reprograming of infected cells (67).

### Inhibition of glutaminolysis significantly reduces MNV replication

Glutaminolysis catabolizes glutamine for anaplerosis and provides a nitrogen source to fuel nucleotide and amino acid biosynthesis, key building blocks required for viral replication (32). The rate-limiting enzyme within the pathway is glutaminase (GLS), which catalyzes the first deaminase reaction (30). Since we uncovered higher glutamine flux in MNV-1 infected cells (Fig. 2), we hypothesized that this pathway would be required for optimal MNV replication. To test this, we infected RAW cells and primary bone marrow-derived macrophages (BMDMs) with MNV-1, CR3, and CR6 at an MOI of 5 for 1 hour. Medium containing CB839, a non-competitive GLS inhibitor, was thus added after infection and infectious titers measured after 8 hours by plaque assay. The concentrations of CB839 used in both RAW cells and primary BMDMs were non-toxic and maintained >80% cell viability (Fig. S1B, C). Cells treated with CB839 had significantly lower MNV titers (by ∼1.5-log_10_) than cells that were treated with vehicle control (Fig. 3A). RAW cells are transformed macrophages, and transformed cells can have altered metabolic processes (80). Thus, to confirm the phenotype observed in RAW cells, we repeated infections in primary BMDMs. MNV-infected primary BMDMs treated with CB839 harbored significantly lower MNV titers (by >1.0-log_10_) than vehicle control (DMSO) cells for all strains despite using a slightly higher non-toxic concentration of CB839 (Fig. 3B). The results in BMDMs confirmed what was seen in RAW cells and showed that glutaminolysis is required for optimal replication of acute and persistent MNV strains.

**Figure 3:**
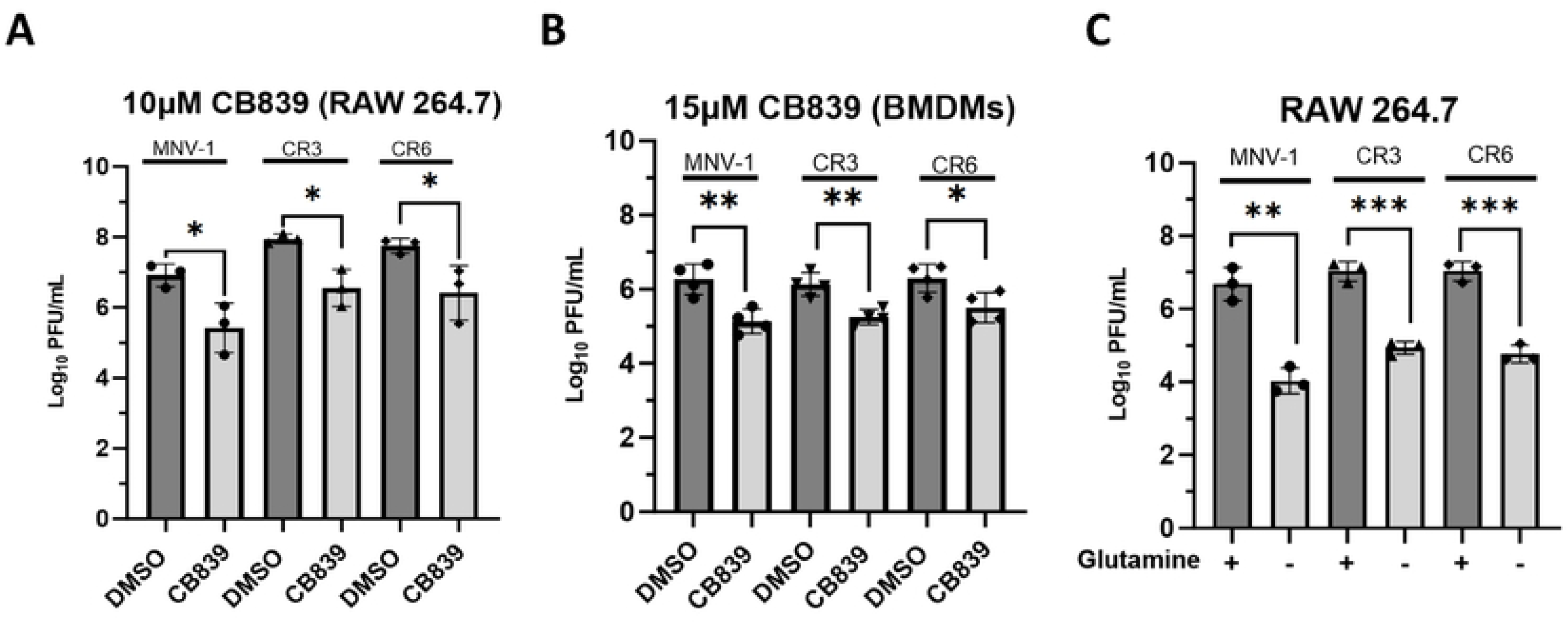
Inhibition of glutaminolysis significantly reduces MNV replication in both primary and transformed macrophages. (**A**) RAW 264.7 cells or (**B**) primary bone marrow-derived macrophages were infected for 1 hour at an MOI of 5 with either MNV-1, CR3, or CR6. Virus inoculum was removed and replaced with medium containing (**A**) 10 μM or (**B**) 15 μM CB839 or vehicle control (DMSO). (**C**) RAW 264.7 cells were infected as before but infection was performed with glutamine-free or replete medium. After an 8 hr incubation, MNV titers were measured via plaque assay. Experiments represent combined data from at least three independent experiments. Statistical analysis was performed using Two-tailed Students-t tests. ***, *P<*0.001; **, *P* <0.01; *, *P<*0.05; ns, not significant.

Pharmacologic inhibitors can result in off-target effects. Hence, we repeated infections in RAW cells with medium lacking glutamine. Infections were performed as before, and viral titers were measured by plaque assay at 8 hpi. While glutamine deprivation has been reported to negatively affect cell viability after 48 hours in numerous cell types (40–41), we confirmed that 8-hour incubation without extracellular glutamine did not negatively affect RAW cell viability (> 80% viability) (Fig. S1D). Glutamine deprivation resulted in significantly lower (by 2-2.5-log_10_) MNV titers for all strains tested (Fig. 3C).

Taken together, these results demonstrate that acute and persistent MNV strains have a similar dependence on glutaminolysis for optimal replication.

### MNV genome replication is the stage in the viral life cycle most dependent upon glutaminolysis

Typical of a positive-sense, single-stranded virus, the MNV life cycle involves the following steps: host cell uptake of viral particles, uncoating of the positive-strand viral RNA (vRNA) genome, direct translation of the positive-sense vRNA to produce nonstructural proteins, and synthesis of viral negative-sense RNA strand for eventual production of new positive-strand vRNA, translation of structural proteins, followed by progeny virion assembly, maturation, and finally egress. To identify the stage within the MNV lifecycle that is most dependent upon glutaminolysis, we continued investigating infection under glutamine-starved conditions to avoid potential off-target effects of CB839. Since glutamine can be used as a nitrogen source for nucleotide biosynthesis (32), we first sought to analyze the role of glutaminolysis on viral genome replication. To test this, RAW cells were infected with MNV-1, CR3, or CR6 for 1 hour at an MOI of 5. After 1 hour, the virus inoculum was replaced with glutamine-free medium, and cells were incubated for 8 hours. After the incubation period, we extracted RNA and assessed viral genome levels via reverse transcriptase quantitative polymerase chain reaction (RT-qPCR). Glutamine-deprived cells had significantly fewer genome copies for all three strains, a 1.8-2.0-log_10_ decrease (Fig. 4A).

**Figure 4:**
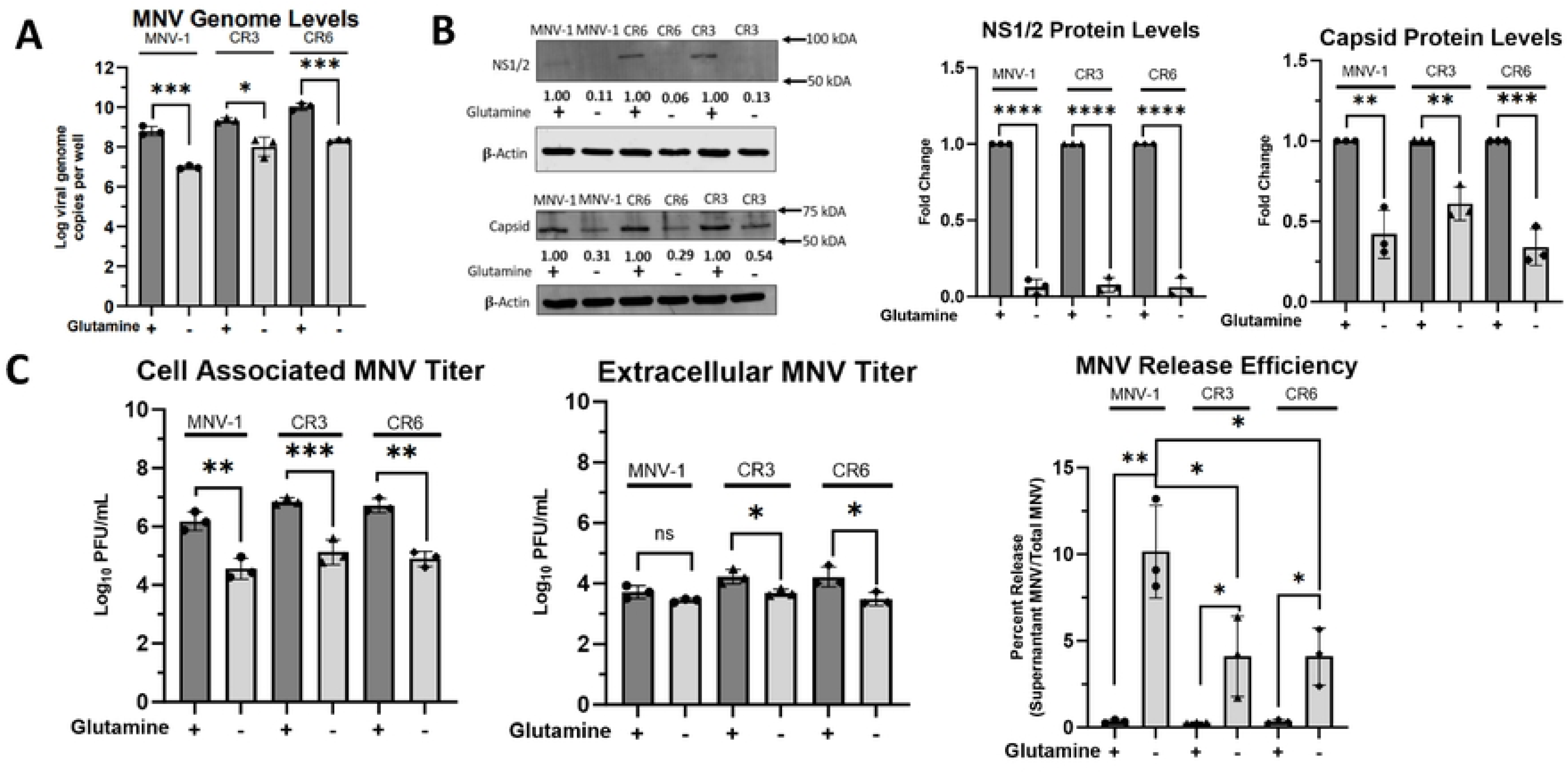
Viral genome replication is the stage of the MNV lifecycle that is most dependent on host glutaminolysis. (**A**) RAW 264.7 cells were infected for 1 hour at an MOI of 5 with either MNV-1, CR3, or CR6. Virus inoculum was removed and replaced with glutamine-free or replete medium. Infected cells were incubated for 8 hours. RNA was extracted and MNV genome levels were assessed via qRT-PCR. (**B**) RAW 264.7 cells were infected as above, and Western blot analysis was performed for MNV viral proteins NS1/2 and capsid. β-actin was used as a loading control. Data shown are representative Western blots from 3 independent experiments. Numbers below blots indicate densitometry measurement of protein level relative to MNV-infected cells receiving replete medium. (**C**) RAW 264.7 cells were infected as before. Supernatants and cell-associated virus were measured separately via plaque assay. Experiments represent combined data from at least three independent experiments. Statistical analysis was performed using Two-tailed Students-t tests and One-Way ANOVA. ***, *P<*0.001; **, *P* <0.01; *, *P<*0.05; ns, not significant.

Glutamine can also be used for amino acid synthesis (81). Thus, we next investigated whether MNV protein synthesis is dependent on host glutaminolysis. RAW cells were infected as before, and after the 8 hr incubation period, levels of the non-structural protein NS1/2 and the capsid protein were measured via western blot (Fig. 4B). NS1/2 protein levels were low in samples from infections with glutamine-containing medium, but not detectable in protein samples from infections with glutamine-free media (Fig. 4B left panel). Quantification of NS1/2 protein signals from three independent replicates indicated a >90% decrease for all strains tested when grown in glutamine-free medium (Fig. 4B middle panel), indicating that glutaminolysis is required for NS1/2 synthesis. Quantification of the capsid protein also showed significantly lower levels of this protein during glutamine starvation (Fig. 4B left panel). For MNV-1 and CR6 infected cells starved for glutamine, we observed a ∼60% reduction in capsid protein levels compared to infections in replete media, while a ∼40% reduction was observed for CR3-infected glutamine-starved cells (Fig. 4B right panel). These data suggested that glutaminolysis is important for MNV viral protein synthesis, although CR3 was slightly more resistant to glutamine starvation than MNV-1 and CR6 (Fig. 4B).

Last, we investigated viral assembly and egress, the end stages of infection. RAW cells were infected with MNV-1, CR3, or CR6 as before in replete and glutamine-starved media. After the 8 hr incubation period, supernatants and cell monolayers were collected separately to measure viral titers and calculate the released virus. In the cell-associated fraction, about a 2.0-log_10_ decrease in viral titers was observed during glutamine starvation vs. replete media for all three strains tested (Fig. 4C left panel), which was similar to the results obtained for total MNV titers (Fig. 3C). The significant decrease in cell-associated MNV titers during glutamine starvation suggests that glutaminolysis is required for MNV assembly in both persistent and acute strains. However, analysis of extracellular MNV showed a significant decrease of MNV titers in glutamine-depleted media only for the persistent strains (Fig. 4C middle panel). Specifically, we observed a 0.75-log_10_ decrease in extracellular CR3 and CR6 titers but no significant decrease for MNV-1 titers (Fig. 4C middle panel), highlighting strain-specific dependencies on glutaminolysis. Additionally, we calculated the ratio of released-to-total viral titers to investigate whether glutamine deprivation affects MNV release efficiency. Surprisingly, glutamine deprivation led to increased release efficiency in all strains, with the highest increase in release efficiency observed in MNV-1 infected cells (Fig. 4C right panel).

In summary, because glutamine can be used for nucleotide synthesis but no change in the intracellular amino acid pool was detected in MNV-1–infected cells in our flux analysis (Fig. S2B), we conclude that genome replication is the stage of the MNV lifecycle that most imminently relies on host glutaminolysis. All other phenotypes observed during later stages of the viral life cycle are most likely a consequence of this initial effect.

### Glutaminase activity is upregulated during MNV infection

Our previous data indicated that glutaminolysis is upregulated during and required for optimal MNV replication. Therefore, we were interested in whether MNV infection increases glutaminolysis through changes in GLS expression. We first directed our attention to GLS transcript and protein levels, since HCMV and HIV have previously been shown to increase GLS protein levels and mRNA expression, respectively (35, 43). To test whether MNV infection modulates GLS expression, we infected RAW cells with MNV-1, CR3, and CR6 for 1 hour at an MOI of 5. After 8 hours, we assessed GLS transcript and protein levels via RT-qPCR and western blot, respectively. We observed that *GLS* transcript levels were significantly higher in MNV-infected cells compared to mock-infected cells (Fig. 5A). Using the housekeeping gene beta-actin as a measure of baseline transcription, we observed some strain-specific differences, with MNV-1 infection leading to a 3-fold increase in *GLS* transcript levels and the persistent strains leading to a 0.5-1-fold increase (Fig. 5A). Western blot analysis of GLS protein levels resulted in no observable difference between MNV and mock-infected cells (Fig. 5B). The two bands present in the immunoblot potentially represent the two isoforms of GLS, KGA and GAC, which are identical in all aspects except the C-terminal domain (45). Surprisingly, quantification of GLS protein levels revealed a small but significant decrease (5-7%) in GLS protein levels in MNV-infected relative to mock-infected cells (Fig. 5B). From these data, we conclude that the upregulation of glutamine metabolism during MNV infection is not due to increased GLS mRNA or protein expression.

**Figure 5:**
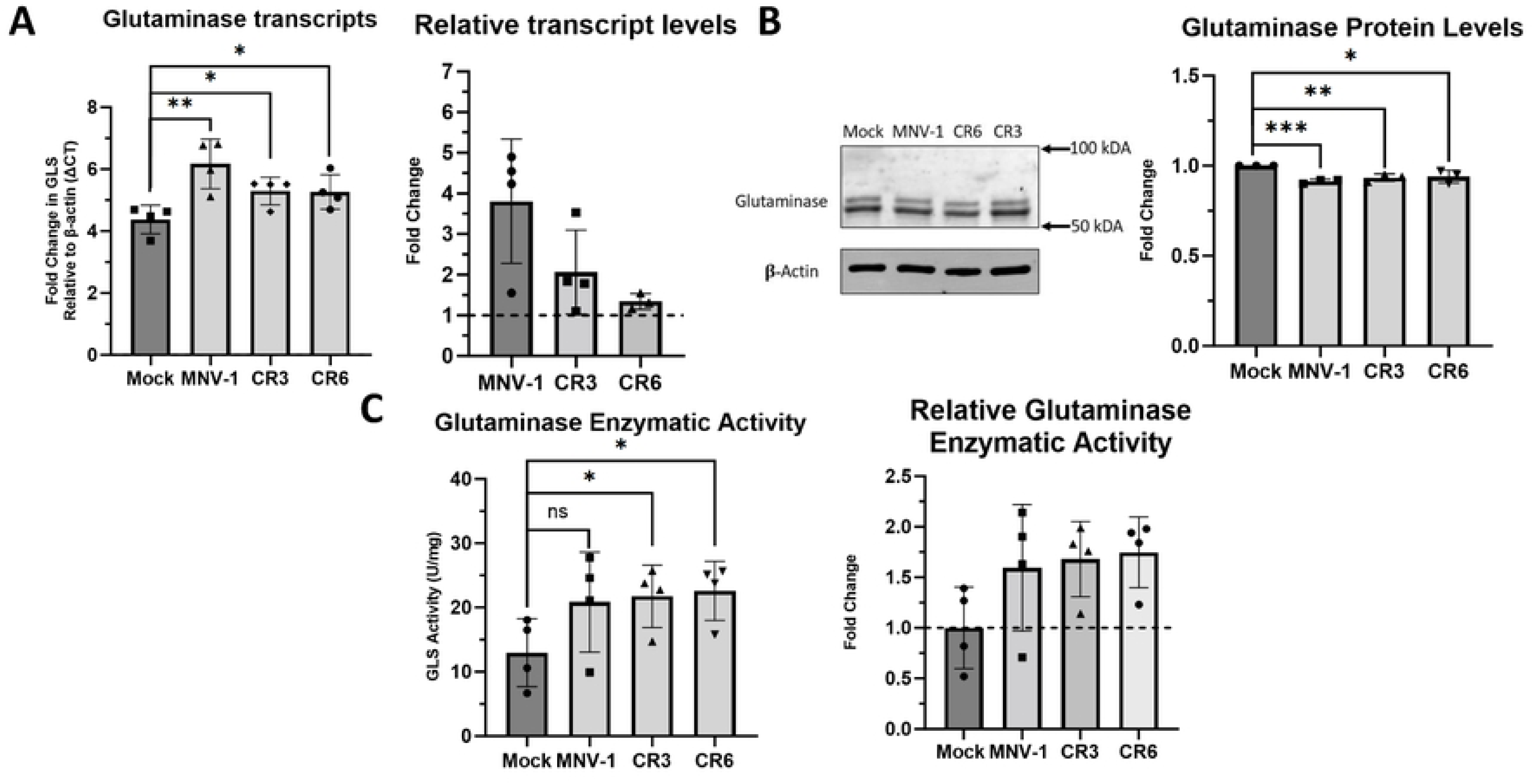
Glutaminase activity is upregulated during MNV infection in macrophages. (A) RAW 264.7 cells were infected for 1 hour at an MOI of 5 with either MNV-1, CR3, or CR6. Virus inoculum was removed and replaced with replete medium. Infected cells were incubated for 8 hours. RNA was extracted and glutaminase transcripts were assessed via qRT-PCR. **(B)** RAW 264.7 cells were infected as before. Western blot analysis was then performed for glutaminase protein levels. β-actin was used as a loading control. A representative Western blot is shown on the left and quantification from 3 independent experiments on the right. **(C)** RAW 264.7 cells were infected as before. Glutaminase activity was analyzed utilizing the Cohesion Biosciences Glutaminase Microassay kit. Experiments represent combined data from at least three independent experiments. Experiments represent combined data from at least three independent experiments. Statistical analysis was performed using Two-tailed Students-t tests. ***, *P<*0.001; **, *P* <0.01; *, *P<*0.05; ns, not significant.

We next investigated whether GLS enzymatic activity was increased during MNV infection, which would be consistent with our flux analysis results showing increased glutamine catabolism during MNV infection. RAW cells were infected with MNV-1, CR3, and CR6 for 8 hours as before and GLS enzymatic activity was analyzed with a commercially available kit that measures ammonia, the byproduct of the reaction that GLS catalyzes (45). We observed higher levels of GLS enzyme activity in MNV-infected cells than in mock-infected cells (Fig. 5C). When analyzing the fold change in GLS activity over mock infected cells, an approximately 0.75-fold increase was detected for all three MNV strains, with each strain increasing GLS activity to a similar extent (Fig. 5C).

Overall, we conclude that increased rates of glutaminolysis during MNV infection in macrophages is the result of increases host cell GLS enzymatic activity, but not due to changes in GLS transcript or protein levels.

### NS1/2 is a viral mediator of increased GLS activity

Viral proteins can mediate changes to host metabolism to ensure optimal infection. For example, dengue virus NS1 interacts with glyceraldehyde-3-phosphate dehydrogenase to upregulate glycolysis (46). Therefore, we investigated whether increased GLS activity in MNV-infected cells is mediated by a viral protein. To test this, we overexpressed individual MNV non-structural proteins in Huh-7 cells expressing the viral receptor CD300lf and measured GLS activity as before. As a control, we first tested whether MNV infection of CD300lf-expressing Huh-7 cells would be sensitive to glutaminolysis inhibition. Cell viability studies determined the concentration of CB839 at which >80% cell viability is maintained to be 5 μM (Fig. S1E). We then infected the cells with MNV-1, CR3, and CR6 for 1 hour at an MOI of 5 before adding medium containing 5 μM CB839 or vehicle control (DMSO) for 8 hrs. Viral titers were measured via plaque assay. We observed a 0.5-1-log_10_ decrease in MNV titers when glutaminolysis was inhibited, confirming that similar to infected macrophages CD300lf-expressing Huh-7 cells are sensitive to glutaminolysis inhibition (Fig. 6A) and provide an efficient cell line for protein overexpression.

**Figure 6:**
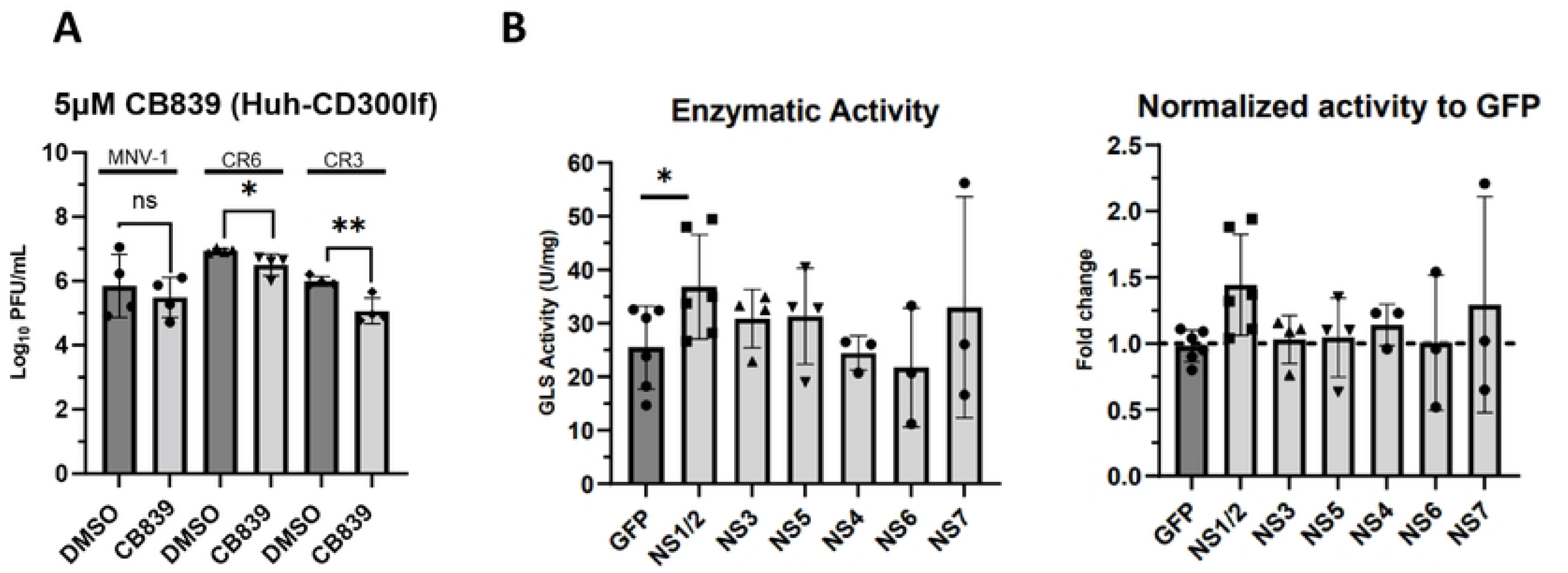
NS1/2 is a viral mediator of increased glutaminase activity in macrophages. (**A**) Huh-7 cells expressing the viral receptor CD300lf were infected for 1 hour at an MOI of 5 with either MNV-1, CR3, or CR6. Virus inoculum was removed and replaced with medium containing 5 μM CB839 or vehicle control (DMSO). Infected cells were incubated for 8 hours and MNV titers were measured via plaque assay. (**B**) Huh-7 CD300lf cells were transfected with plasmids encoding the indicated MNV-1 non-structural protein or green fluorescent protein. Transfected cells were incubated for 24-48 hours. Glutaminase activity was analyzed utilizing the Cohesion Biosciences Glutaminase Microassay kit. Experiments represent combined data from at least three independent experiments. Statistical analysis was performed using Two-tailed Students-t tests. **, *P* <0.01; *, *P<*0.05; ns, not significant.

Having confirmed the importance of glutaminolysis during MNV infection in CD300lf-expressing Huh-7 cells, we investigated whether the expression of an individual viral protein would alter GLS activity. To this end, we transfected CD300lf expressing Huh-7 cells with plasmids for the expression of 6 MNV-1 non-structural proteins (NS1/2 and NS3 to NS7) or green fluorescent protein (GFP) as a negative control. Transfected cells were incubated for 24-48 hours, and cell lysates were first tested for successful protein expression via western blot (Fig. S3). After confirming expression of the proteins of interest, cell lysates were analyzed for GLS activity. We observed increased GLS activity in cells expressing NS1/2 (Fig. 6B left panel), with an approximately 0.5-fold change in GLS activity (Fig. 6B, right panel), over cells expressing GFP. This is slightly less than the 0.75-fold increase in GLS activity observed during MNV-1 infection (Fig. 5C). NS7 overexpression resulted in highly variable GLS activities but was not statistically significant (Fig. 6B). Thus, other viral proteins, e.g. NS7 or structural proteins, may contribute to the full increase in GLS activity observed in MNV-infected macrophages.

Taken together, these data demonstrate that the MNV structural protein NS1/2 mediates an increase in GLS activity and is a viral factor upregulating glutaminolysis during macrophage infection.

## Discussion

Viruses have evolved numerous mechanisms for manipulating host cellular metabolism to create a more favorable intracellular environment to support optimal replication. Our previous study showed that MNV-1 infection significantly alters numerous host metabolic pathways, including glycolysis, the PPP, and OXHPOS, thereby supporting the energetic and biosynthetic needs for optimal virion production (29). In our present study, we extended our investigation to include two persistent MNV strains, CR3 and CR6, and observed strain-dependent differences compared to MNV-1 in that while these strains also required host glycolysis and the PPP for optimal replication, they did not require OXPHOS. To support the previous static metabolomic analysis, we furthermore performed metabolic flux analysis to measure the incorporation of labeled carbon from glucose and glutamine. These data showed significantly higher glucose and glutamine catabolism during MNV-1 infection, thus supporting the observation that MNV infection upregulates both metabolic pathways concurrently. Having previously investigated the role of glycolysis during MNV infection, we focused on the role of glutaminolysis during MNV infection in this study. Glutamine deprivation and pharmacological inhibition of glutamine catabolism resulted in significantly lower MNV-1, CR3, and CR6 viral titers in multiple cell types, thus revealing that glutaminolysis is required for optimal MNV replication. Our results also showed that MNV genome replication is the first step in the viral life cycle that depends on glutaminolysis and our mechanistic studies point to NS1/2 as a viral protein that mediates upregulation of GLS activity, the key enzyme in glutaminolysis. Thus, in addition to glycolysis, glutaminolysis is another intrinsic host metabolic factor that contributes to optimal MNV replication. Collectively, our investigation has revealed both shared and strain-specific metabolic dependencies that may underly the different pathogenic phenotypes of various MNV strains.

Glycolysis and glutaminolysis are the catabolic pathways for glucose and glutamine, respectively, and these molecules are the main carbon sources used by mammalian cells to perform a myriad of cellular processes. Importantly, these pathways are often concurrently rewired by viruses, since metabolites from the glycolytic pathway can not only be used for energy production via OXPHOS, but also can be used within the PPP for nucleotide synthesis, molecules that viruses need for genome replication. Additionally, glycolytic intermediates can be used in lipid biosynthesis, and when glycolytic intermediates are used more for lipid biosynthesis or lactic acid assembly rather than energy production, aKG, a glutaminolysis product, can be shuttled into the TCA cycle to ensure continuous downstream ATP production via anaplerosis. This phenotype is observed in HCMV infections (35). Glutamine catabolism also provides nitrogen-containing metabolites for amino acid and nucleotide biosynthesis (32). Together, glycolysis and glutaminolysis provide the necessary building blocks and energetic needs for optimal progeny virion production. Hence, viruses may target both pathways to promote optimal replication. In the present study we observed increased glycolysis and glutaminolysis during MNV-1 replication in murine macrophages. Glutamine deprivation and treatment with the pharmacological inhibitor CB839 significantly decreased virion production of MNV-1, CR3, and CR6 through reduced genome replication, which resulted in lower levels of non-structural and structural protein synthesis, viral assembly, and release. Diverse viruses such as HIV-1, white-spot syndrome virus, hepatitis C virus, influenza virus, and adenovirus also upregulate both glycolysis and glutaminolysis during infection (9, 47–55). However, the molecular mechanisms underlying the upregulation of these two key metabolic pathways and how this metabolic rewiring affects virus replication vary by virus and host cell type. Uncovering these mechanisms may reveal shared metabolic dependencies and therapeutic chokepoints.

As obligate intracellular parasites, viruses rely on the metabolic products of host cells and have evolved capabilities to hijack metabolic resources and stimulate specific metabolic pathways required for replication. However, the viral proteins responsible for metabolic control are mostly unknown. In this study, we identified the NoV non-structural protein NS1/2 as being involved in host cell metabolic modulation. This protein is released from the viral polyprotein precursor via proteolytic activity of the viral protease NS6 (82). Our results strongly suggest that upon release from the polyprotein one function of the NS1/2 protein is to enhance GLS enzymatic activity, leading to increased glutaminolysis. Other viral proteins known to mediate changes to host metabolism come from diverse virus families. For example, three different DNA viruses use non-structural proteins to modulate host metabolism. Epstein-Barr virus increases fatty acid synthase expression during lytic replication through the immediate-early non-structural protein BRLF1, which works in a p38 stress mitogen-activated protein kinase-dependent manner to increase fatty acid production (56). Hepatitis B virus uses viral protein X to reprogram liver glucose metabolism through increased expression of key gluconeogenic enzymes (57). And adenoviruses use the E4ORF1 gene product through a direct interaction with c-Myc to increase anabolic glucose metabolism and glutaminolysis (9,49). Enterovirus A71, on the other hand, affects host cell metabolism through its structural protein VP1, which directly binds to trifunctional carbamoyl-phosphate synthetase 2, aspartate transcarbamylase, and dihydroorotase to promote increased pyrimidine synthesis (37). These examples highlight that both non-structural and structural viral proteins from diverse viral families can contribute to altering host metabolism during viral infection. However, our work on NS1/2 increasing GLS activity provides the first example of an RNA virus that upregulates glutaminolysis through a specific non-structural viral protein. Although we cannot rule out that NS1/2 is the only MNV viral protein that increases GLS activity. Future investigations into the detailed mechanism of NS1/2-mediated increase in GLS enzymatic activity are needed and have the potential to reveal fundamental insights into norovirus-host interactions and pathogenesis.

Macrophages are highly plastic immune cells that adapt to different physiological microenvironments. These cells are often parsed into two major categories: pro-inflammatory (M1) and anti-inflammatory/pro-resolving (M2) macrophages (58). Importantly, these two macrophage phenotypes are associated with distinct metabolic profiles. Hallmarks of M1 macrophages include high rates of glycolysis, fatty acid synthesis, and pentose phosphate activity. In contrast, hallmarks of M2 macrophages include high rates of glutaminolysis, fatty acid oxidation, and OXPHOS (58). Our previous (29) and current metabolomic analyses revealed significant upregulation of central carbon metabolism and increased carbon flow through glycolysis and glutaminolysis during MNV infection of macrophages. Since upregulation of glycolysis, the PPP, and increased succinate production are hallmarks of M1 macrophages, while upregulation of glutaminolysis and OXPHOS are hallmarks of an M2 macrophage, MNV-infected macrophages display a hybrid metabolic profile during infection. Intriguingly, the underlying metabolic program is crucial for macrophage function (58). However, how the metabolic alterations induced by MNV infection impact macrophage function remains unknown. Like MNV, bacteria also rewire macrophage metabolism to grow and evade innate immunity. *Legionella pneumophila, Brucella abortus,* and *Listeria monocytogenes* rewire macrophages towards aerobic glycolysis, and *L. pneumophila* enhances glycolysis by a yet-to-be-determined mechanism (59). *L. monocytogenes* uses a bacterial toxin to induce mitochondrial fragmentation and takes advantage of increased glycolysis in M1 macrophages to efficiently proliferate (60). While chronic *B. abortus* infection preferentially occurs in M2 macrophages, it requires PPARγ to increase glucose availability (61). Parasites can also alter macrophage metabolism during intracellular infection. For example, *Leishmania spp*. are protozoan parasites that infect macrophages and activate HIF-1α to upregulate HIF-1α target genes, including glucose transporters and glycolytic enzymes, resulting in increased glucose uptake, glycolysis, and activation of the PPP (62). These examples suggest that while MNV infection increases the availability of resources for optimal infection, rewired macrophage metabolism may also promote changes to the host immune response. Disentangling which metabolic pathways are directly altered by MNV and which are consequences of macrophage host defenses is an important area for future investigations.

In conclusion, we have shown that glutaminolysis, in addition to glycolysis, is an intrinsic host factor promoting optimal replication of MNV. Our data are consistent with a model whereby MNV uses the NS1/2 protein to upregulate GLS activity during infection of macrophages, which increases glutamine catabolism. Our previous and current findings reveal that central carbon metabolism plays an important role in NoV replication, and these findings may uncover novel chokepoints for therapeutic intervention and new avenues for improving HNoV cultivation.

## Methods

### Compounds and reagents

2-Deoxyglucose (2DG) (Sigma #D8375) was solubilized fresh for each experiment in cell culture medium to 100 mM and added to the culture medium at a final concentration of 10 mM. CB839 (Cayman Chemical #22038) was solubilized in DMSO at 10 mM and used at final concentrations of 5, 10, or 15 μM. 6-Aminonicotinamide (6AN) (Cayman #10009315) was solubilized in DMSO at 500 mM and used at 500 or 750 μM. Oligomycin A (Cayman #11342) was solubilized in DMSO at 5 mM and used at 1 μM. Glutamine-free media was prepared fresh for each experiment using DMEM-10 medium (Gibco DMEM medium #11995-044 with 4.5 g/L D-Glucose, 10% dialyzed fetal bovine serum (Thermo Fischer Scientific #A3382001), and 1% HEPES buffer (1M, Gibco #15630-080). MNV-1 NS1/2, NS3, and NS5 plasmids were a kind gift from Dr. Jason Mackenzie (University of Melbourne, AUS) and previously described (83). Flag-tagged MNV-1 NS4, NS6, and NS7 plasmids were a kind gift from Dr. Ian Goodfellow (University of Cambridge, UK) and previously described (84).

### Cell culture and virus strains

The RAW 264.7 macrophage cell line (referred to herein as RAW cells) (ATCC TIB-71) and CD300lf-expressing Huh-7 cells were maintained in DMEM-10 medium (Gibco DMEM medium #11995-065 with 4.5 g/L D-Glucose and 110 mg/L Sodium Pyruvate, 10% Fetal Bovine Serum [HyClone #SH30396.03], 1% HEPES buffer [1M, Gibco #15630-080], 1% Non-Essential Amino Acids [100X, Gibco #11140-050] and 1% L-Glutamine [200 mM, Gibco #25030-081]) in treated tissue culture flasks at 37°C/5% CO_2_. CD300lf-expressing Huh-7 cells were a gift from Dr. Stefan Taube (University of Lübeck, Germany) and were previously described (63). Primary bone marrow-derived macrophages (BMDM) were differentiated from male Balb/C mouse femur and tibia bone marrow in 20% L929 medium (Gibco DMEM medium, 20% FBS [HyClone #SH30396.03], 30% L9 supernatant, 1% L-Glutamine, 1% Sodium Pyruvate, 0.25 mL β-mercaptoethanol/L and 2% Penicillin/Streptomycin). All experiments using primary cells were performed with 10% L929 working medium (same as 20% L929 medium but with 10% L929 supernatant). The plaque purified MNV-1 clone (2002/USA) MNV-1.CW3 (referred herein as MNV-1) was used at passage 6 in all experiments. CR3 and CR6 were also used at passage 6 in all experiments (64).

### Virus infections and plaque assay

All MNV infections were performed in the RAW 264.7 cell line, Balb/C primary bone marrow-derived macrophages (BMDM), or CD300lf-expressing Huh-7 cells. Cells were grown in 12-well tissue culture plates seeded at 5×10^5^ cells/well. At the time of infection, the medium was replaced with 1 mL of media containing MNV-1, CR3, or CR6 at the indicated MOI. Plates were rocked for 1 hour on ice. Then, cells were washed 3X with cold DPBS++ (+Calcium and +Magnesium Chloride—Gibco #14040), fresh medium was added containing metabolic inhibitors at the indicated concentrations, vehicle control, or glutamine-free media. Cells were incubated for indicated times. Cells were then frozen at −80°C and freeze-thawed two times before lysates were analyzed by plaque assay as previously described (65). Vehicle control experiments were performed using DMSO in a v/v match to the volume of metabolic inhibitors. Primary cell infections were done the same as RAW infections except in medium containing 10% L929 supernatant.

### RNA extraction and RT-qPCR

Experiments to quantify MNV genome copies and glutaminase expression were performed on MNV- or mock-infected RAW cells as indicated above. At time of RNA extraction, cells were washed 1X with cold DPBS++ and then 500 μL of Zymo Research TriReagent (#R2050-1) was added. Extraction was performed per manufacturer’s directions using the Zymo Research Direct-zol RNA MiniPrep Plus (#R2072) and then used for One-Step TaqMan Assay. Primers used to measure murine glutaminase transcript and MNV genome levels were previously described (66, 78).

### Protein extraction, SDS-PAGE, and immunoblotting

Experiments were performed as described above in 12-well or 6-well tissue culture plates. At time of harvest, cells were washed 2X with cold DPBS++ and RIPA buffer (Pierce #89900) containing complete EDTA-free protease inhibitor cocktail (Roche #11873580001) was added to wells. Cells were scraped, moved to Eppendorf tubes, and incubated on ice for 15 minutes. Cells were then spun at 4°C at 14,000 x g for 15 minutes. Lysates were moved to fresh tubes, and Laemmli buffer with β-mercaptoethanol was added at 3:1 lysate to buffer ratio before freezing the sample until analysis. SDS-PAGE was performed with BioRad 4-20% Mini-Protean TGX gels (BioRad #456-1096) per standard SDS-PAGE procedures (79). Gels were transferred to Immobilon-FL transfer membranes (#IPFL00010, pore size 0.45 μm) using a Semi-Dry transfer at 10V for 60 minutes. Membranes were blocked in PBS+0.05% Tween + 1% low-fat milk for 1 hour at room temp, then primary antibodies were added in the same buffer and membranes were rocked at 4°C overnight. Membranes were washed 3X with 1X PBS, then secondary LI-COR fluorescent antibodies were added for 1 hour at room temp and then visualized on the LI-COR Odyssey Imager. Western blots were quantified by densitometry using ImageJ and normalizing bands to β-actin. Antibodies used: mouse mAb β**–**Actin (clone 8H10D10, Cell Signaling #3700) at 1:10,000 dilution; rabbit mAb β-Actin (clone 13E5, Cell Signaling #8457) at 1:10,000 dilution; anti-rabbit polyclonal glutaminase (Proteintech #12855-1-AP) at 1:1000 dilution; anti-mouse monoclonal FLAG (Sigma #F1804) at 1:3000 dilution. The rabbit polyclonal anti-MNV-1 capsid antibody (used at 1:500 dilution) was described previously (29). The mouse monoclonal anti-NS1/2 and anti-NS5 antibodies (both used at 1:3000 dilution) were a kind gift from Dr. Vernon Ward (University of Otago, New Zealand) and previously described (85).

### Cell Viability Assay

Cell viability was tested with the WST-1 Cell Proliferation Reagent (Sigma #5015944001) or Resazurin Cell Viability Assay Kit (Biotium #30025-1). Briefly, RAW cells, primary BMDMS, or CD300lf-expressing Huh-7 cells were plated at 2×10^5^ per well of a 24-well plate. After overnight growth at 37°C/5% CO_2_, medium was replaced with DMEM-10 medium containing a specific pharmacological inhibitor. Treated cells were then placed back at 37°C/5% CO_2_ for a 24-hour incubation period. The following day, cell viability was calculated according to the manufacturer’s recommendations.

To measure the viability of RAW cells in glutamine-free media, cells were plated at 5*10^5 per well in a 6-well plate. After overnight growth at 37°C/5% CO_2_, media was replaced with glutamine-free DMEM-10 medium for 8 hours. After the incubation, cells were scrapped with a cell scrapper and cell viability was measured using trypan blue staining on a Life Technologies Countess 3 automated cell counter assay platform. Cell viability was calculated as the percent of live cells in glutamine-free media treated vs. untreated controls.

### Metabolic Flux Analysis

5×10^5^ RAW cells were plated in 6-well plates and infected with MNV-1 or mock-infected as described above. Following the removal of the virus inoculum, fresh medium was added containing uniformly labeled ^13^C_5_ glucose or glutamine and incubated at 37°C/5% CO_2_ for 8 hours. Following the 8-hour incubation, cells were washed 2x DPBS (+Calcium and +Magnesium Chloride –Gibco #14040) and 300 μL of ice-cold methanol was added. Wells were scraped with a cell lifter and the volume was transferred to a fresh Eppendorf tube where 300 μL of water containing 1µg of norvaline internal standard was added to each tube. Next, 600µL of high-performance liquid-chromatography grade chloroform was added to each tube to isolate nonpolar lipid content from the sample matrix. Tubes were then vortexed at 4°C for 30 minutes and centrifuged at 17,000 x g for 15 minutes at 4°C to separate contents into an upper aqueous layer and lower chloroform layer. The upper phase was collected into new tubes which were then dried by vacuum centrifugation in a SpeedVac for 5 hours at room temperature. After drying, samples were stored at −80°C until GC-MS analysis. For polar metabolite analysis, dried samples were derivatized with 30µL of 2% methoxyamine hydrochloride in pyridine at 45°C for 1 hour under constant shaking. Then 30 µL of N-tert-butyldimethylsilyl-N-methyltrifluoroacetamide (MBTSTFA) + 1% tertbutyldimetheylchlorosilane (TBDMCS) was added, and samples were further incubated at 45°C for 30 min. Derivatized samples were then transferred to GC vials with glass inserts and loaded for autosampler injection. GC-MS analysis was performed using an Agilent 7890 GC equipped with a 30m DB-35MS UI capillary column connected to an Agilent 5977B MS. Samples were run with 1 mL/min helium flow with the following heating cycle for the GC oven: 100 °C for 1 minute, ramp of 3.5 °C/min to 255 °C, ramp of 15 °C to 320 °C, then held at 320 °C for 3 min to a total run time of 52.6 min. MS source was held at 230 °C and quadrupole at 150 °C. Data was acquired in scan mode (70-600 m/z). The relative abundance of metabolites was calculated from the integrated signal of all potentially labeled ions for each metabolite fragment. Metabolite levels were normalized to the norvaline internal standard and quantified using 10-point calibration with external standards for 36 polar metabolites. Mass Isotopomer Distributions (MIDs) were corrected for natural isotope abundances and tracer purity using IsoCor.

### Overexpression of Viral Proteins

A total of 2.0 μg of plasmid DNA harboring sequences for individual MNV non-structural proteins or green fluorescent protein (GFP) was added to 100 μL of Opti-MEM media (Thermo Fischer Scientific #11058-02 with L-Glutamine and HEPES). Then, 8 μL of FuGENE HD Transfection reagent (FuGENE #0000553572) was added to the Opti-MEM plasmid mix and centrifuged for 10 s at 8000 x g. Plasmid mix was then incubated for 15 minutes at room temperature. After the incubation, the plasmid mix was added to a separate Eppendorf tube containing 1.6×10^6^ CD300lf-expressing Huh-7 cells and incubated for 10 minutes at room temperature. After the incubation, 500 μL of the cell suspension was plated per well in a 6-well plate and incubated at 37°C/5% CO_2_ for 24-48 hours. After the incubation period, two of the wells were used to confirm successful expression of the viral protein via western blot analysis as described above. The remaining well was used to analyze glutaminase activity with the commercially available Cohesion Biosciences Microplate Assay Kit as described above.

### Glutaminase Activity Assay

Glutaminase enzymatic activity was assessed with the commercially available Cohesion Biosciences Microplate Assay Kit (#CAK1065). Briefly, RAW or CD300lf-expressing Huh-7 cells were either mock- or MNV-infected as described above. After 8 hours of incubation at 37°C/5% CO_2_, cells were sonicated for 10 seconds 30x and kit contents added per the manufacturer’s instructions. Samples were transferred to a 96-wellplate and absorbance at 620 nm was measured in a Synergy H1 plate reader. Glutaminase activity was calculated following the manufacturer’s instructions.

### Statistical Analysis

For all experiments, data were analyzed in Prism9 using the tests as indicated in figure legends.

## Acknowledgements

These studies were funded by the University of Michigan Pandemic Relief fund to C.E.W. D.N. and N.M. are supported by NCI grant nos. R01CA227622 and R01CA204969. D.N. is also supported by grants from the Rogel Cancer Center and the Forbes Institute for Cancer Discovery. A.H. was supported by the Molecular Mechanisms of Microbial Pathogenesis Training Grant (5T32AI007528-24). We thank past and present members of the Wobus lab for helpful discussions, and Drs. Ian Goodfellow, Vernon Ward, Jason Mackenzie, and Stefan Taube for the indicated reagents.

## Supplemental figure legends

**Supplementary Figure 1:**
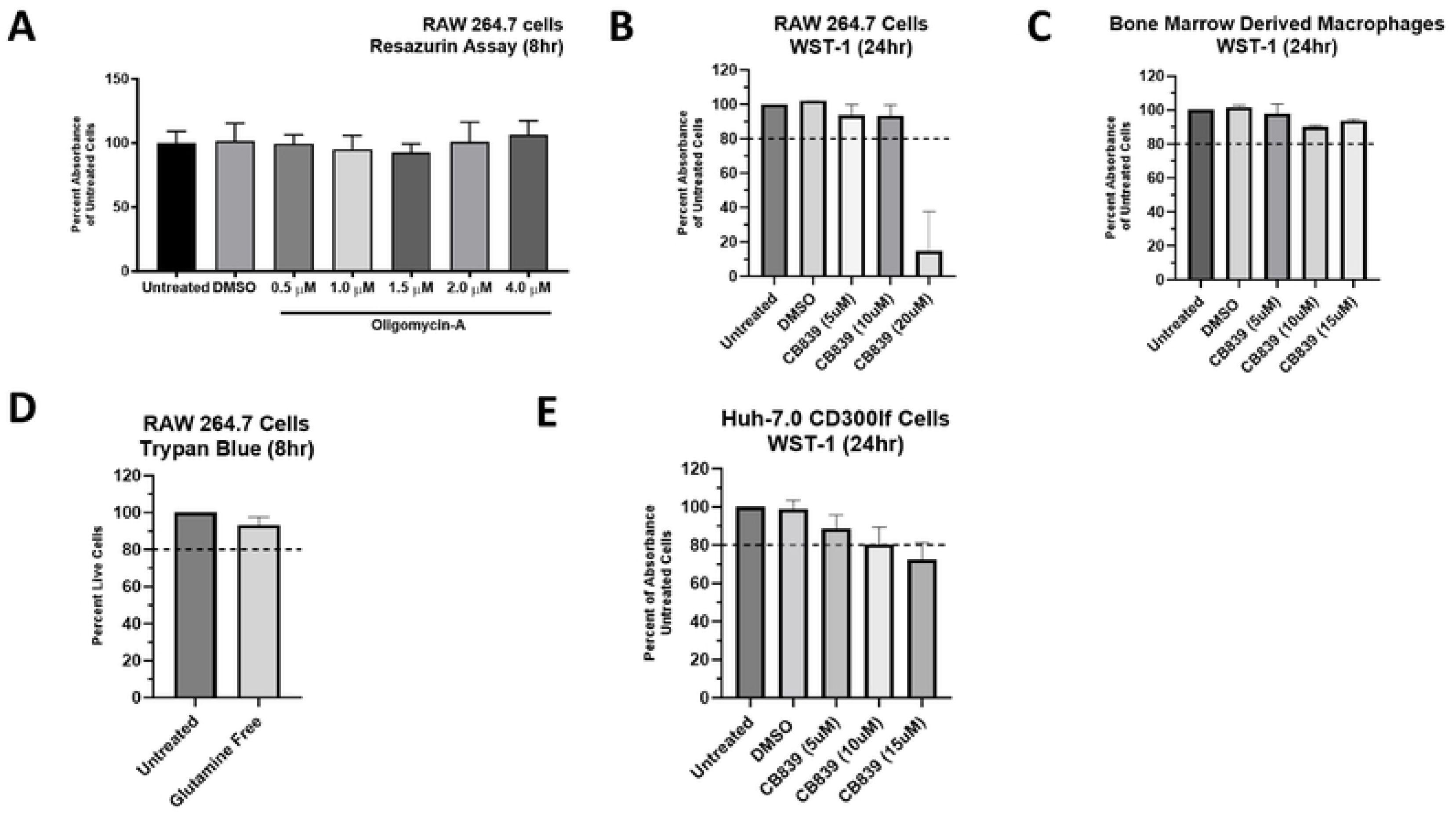
Cell viability assays of indicated cell lines. **(A-B)** RAW 264.7 cells were treated with indicated concentrations of **(A)** Oligomycin-A, **(B)** CB839, or vehicle control (DMSO) for either 8 or 24 hours, respectively. Cell viability was measured using Resazurin or WST-1 reagent. **(C)** Primary bone marrow-derived macrophages were treated with CB839 or vehicle control at the indicated concentrations for 24 hours. Cell viability was measured using WST-1 reagent. **(D)** RAW 264.7 cells were incubated with glutamine free or replete medium for 8 hours. Cell viability was measured using trypan blue staining on a Life Technologies Countess 3 automated cell counter assay platform. **(E)** Huh-7 CD300lf cells were treated with indicated concentrations of CB839 for 24 hrs. Cell viability was measured using WST-1 reagent. Experiments represent combined data from at least two independent experiments with two technical replicates each.

**Supplementary figure 2:**
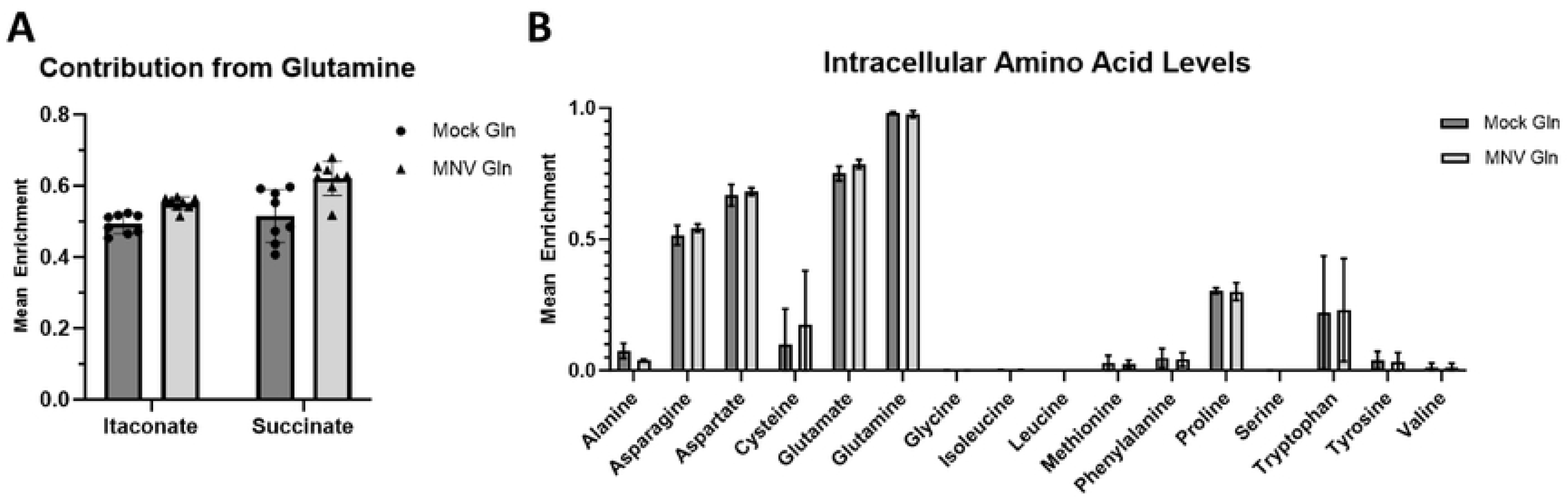
MNV-1 infection does not alter the intracellular amino acid pool. **(A-B)** RAW 264.7 cells were either mock-infected or infected with MNV-1 for 1 hour at an MOI of 5. The virus inoculum was removed and replaced with medium containing ^13^C_5_-glutamine for 8 hours. Intracellular metabolites and amino acids were extracted with ice-cold methanol and measured by mass spectrometry. Experiments represent combined data from two independent experiments with four technical repeats.

**Supplementary figure 3:**
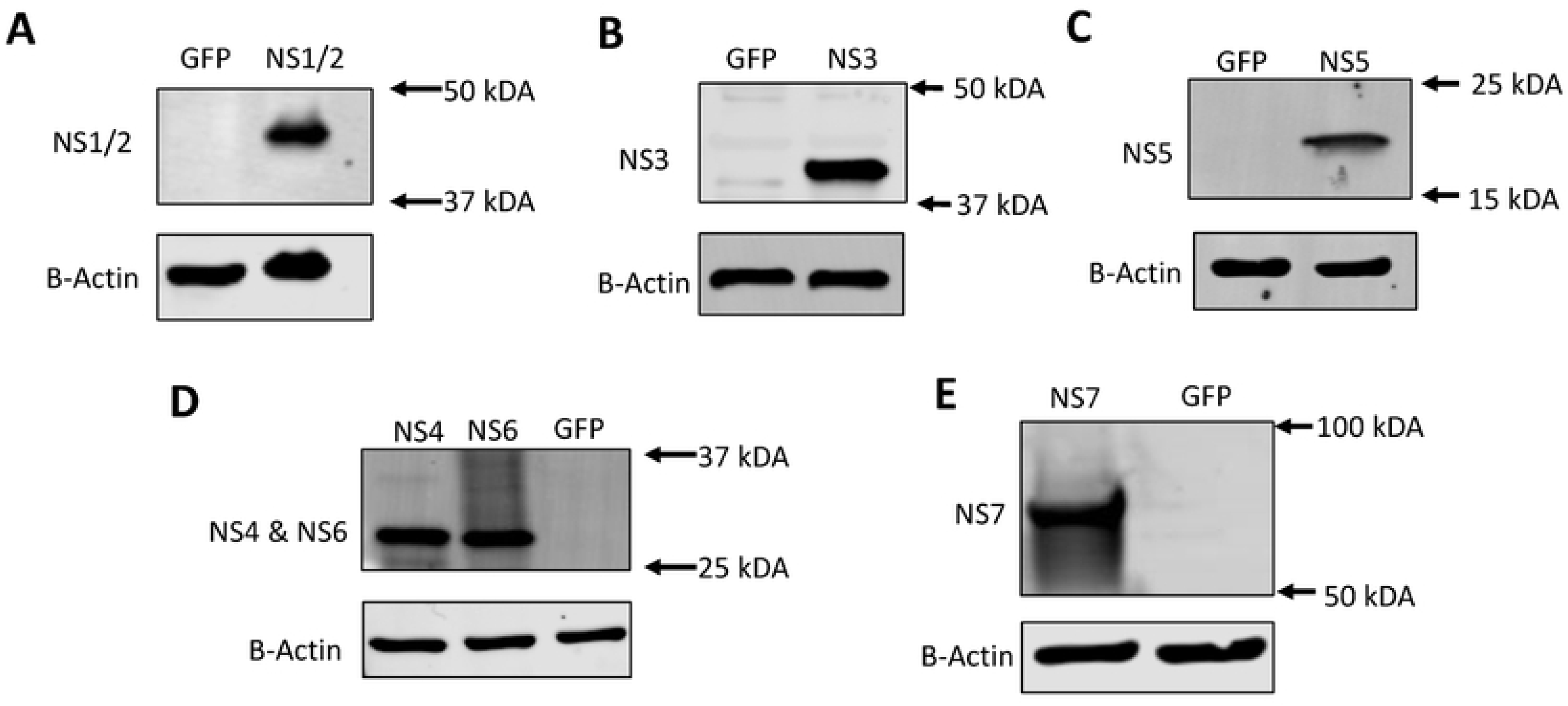
Validation of MNV-1 non-structural protein expression. **Successful expression of MNV viral proteins.** Validation of MNV-1 nonstructural protein expression. (**A-E**) Huh-7 CD300lf cells were transfected with plasmids encoding the indicated MNV-1 nonstructural protein or green fluorescent protein. Transfected cells were incubated for 24-48 hours. Western blot analysis was performed to confirm successful expression. β-actin was used as a loading control. Data shows representative Western blots from 3 independent experiments.

## References

1. Sumbria D, Berber E, Mathayan M, Rouse BT. Virus Infections and Host Metabolism-Can We Manage the Interactions? Front Immunol. 2021 Feb 3;11:594963.

2. Olenchock BA, Rathmell JC, Vander Heiden MG. Biochemical Underpinnings of Immune Cell Metabolic Phenotypes. Immunity. 2017 May 16;46(5):703–713.

3. Kaur J, Debnath J. Autophagy at the crossroads of catabolism and anabolism. Nat Rev Mol Cell Biol. 2015 Aug;16(8):461–72.

4. Goodwin CM, Xu S, Munger J. Stealing the Keys to the Kitchen: Viral Manipulation of the Host Cell Metabolic Network. Trends Microbiol. 2015 Dec;23(12):789–798.

5. Caradonna KL, Engel JC, Jacobi D, Lee CH, Burleigh BA. Host metabolism regulates intracellular growth of Trypanosoma cruzi. Cell Host Microbe. 2013 Jan 16;13(1):108–17.

6. Bravo-Santano N, Ellis JK, Mateos LM, Calle Y, Keun HC, Behrends V, Letek M. 2018. Intracellular Staphylococcus aureus modulates host central carbon metabolism to activate autophagy. mSphere 3:e00174–18

7. Thaker SK, Ch’ng J, Christofk HR. Viral hijacking of cellular metabolism. BMC Biol. 2019 Jul 18;17(1):59

8. Fontaine KA, Sanchez EL, Camarda R, Lagunoff M. Dengue virus induces and requires glycolysis for optimal replication. J Virol. 2015 Feb;89(4):2358–66.

9. Fontaine KA, Camarda R, Lagunoff M. Vaccinia virus requires glutamine but not glucose for efficient replication. J Virol. 2014 Apr;88(8):4366–74.

10. Thaker SK, Chapa T, Garcia G Jr, Gong D, Schmid EW, Arumugaswami V, Sun R, Christofk HR. Differential Metabolic Reprogramming by Zika Virus Promotes Cell Death in Human versus Mosquito Cells. Cell Metab. 2019 May 7;29(5):1206–1216.e4.

11. Shrinet J, Shastri JS, Gaind R, Bhavesh NS, Sunil S. Serum metabolomics analysis of patients with chikungunya and dengue mono/co-infections reveals distinct metabolite signatures in the three disease conditions. Sci Rep. 2016 Nov 15;6:36833

12. Thai M, Graham NA, Braas D, Nehil M, Komisopoulou E, Kurdistani SK, McCormick F, Graeber TG, Christofk HR. Adenovirus E4ORF1-induced MYC activation promotes host cell anabolic glucose metabolism and virus replication. Cell Metab. 2014 Apr 1;19(4):694–701.

13. Chambers JW, Maguire TG, Alwine JC. Glutamine metabolism is essential for human cytomegalovirus infection. J Virol. 2010 Feb;84(4):1867–73.

14. Syed GH, Amako Y, Siddiqui A. Hepatitis B virus hijacks host lipid metabolism. Trends Endocrinol Metab. 2010 Jan;21(1):33–40.

15. Sanchez EL, Lagunoff M. Viral activation of cellular metabolism. Virology. 2015 May;479-480:609–18.

16. Carvajal, J. J., Avellaneda, A. M., Escobar, D., Covián, C., Kalergis, A. M., & Lay, M. K. (2019). Human Norovirus Proteins: Implications in the Replicative Cycle, Pathogenesis, and the Host Immune Response. Frontiers in Immunol. 2020 Jun 16;11:961.

17. Ahmed SM, Hall AJ, Robinson AE, Verhoef L, Premkumar P, Parashar UD, Koopmans M, Lopman BA. Global prevalence of norovirus in cases of gastroenteritis: a systematic review and meta-analysis. Lancet Infect Dis. 2014 Aug;14(8):725–730.

18. Bartsch SM, Lopman BA, Ozawa S, Hall AJ, Lee BY. Global Economic Burden of Norovirus Gastroenteritis. PLoS One. 2016 Apr 26;11(4):e0151219.

19. de Graaf M, van Beek J, Koopmans MP. Human norovirus transmission and evolution in a changing world. Nat Rev Microbiol. 2016 Jul;14(7):421–33.

20. Scallan E, Hoekstra RM, Angulo FJ, Tauxe RV, Widdowson MA, Roy SL et al. Foodborne illnessacquired in the United States--major pathogens. Emerging infectious diseases 2011; 17(1): 7–15.

21. Netzler NE, Enosi Tuipulotu D, White PA. Norovirus antivirals: Where are we now? Med Res Rev. 2019 May;39(3):860–886.

22. Ettayebi K, Crawford SE, Murakami K, Broughman JR, Karandikar U, Tenge VR, Neill FH, Blutt SE, Zeng XL, Qu L, Kou B, Opekun AR, Burrin D, Graham DY, Ramani S, Atmar RL, Estes MK. Replication of human noroviruses in stem cell-derived human enteroids. Science. 2016 Sep 23;353(6306):1387–1393.

23. Estes MK, Ettayebi K, Tenge VR, Murakami K, Karandikar U, Lin SC, Ayyar BV, Cortes-Penfield NW, Haga K, Neill FH, Opekun AR, Broughman JR, Zeng XL, Blutt SE, Crawford SE, Ramani S, Graham DY, Atmar RL. Human Norovirus Cultivation in Nontransformed Stem Cell-Derived Human Intestinal Enteroid Cultures: Success and Challenges. Viruses. 2019 Jul 11;11(7):638.

24. Mirabelli C, Jones MK, Young VL, Kolawole AO, Owusu I, Shan M, Abuaita B, Turula H, Trevino JG, Grigorova I, Lundy SK, Lyssiotis CA, Ward VK, Karst SM, Wobus CE. Human Norovirus Triggers Primary B Cell Immune Activation *In Vitro*. mBio. 2022 Apr 26;13(2):e0017522.

25. Wobus CE, Thackray LB, Virgin HW 4th. Murine norovirus: a model system to study norovirus biology and pathogenesis. J Virol. 2006 Jun;80(11):5104–12.

26. Thackray LB, Wobus CE, Chachu KA, Liu B, Alegre ER, Henderson KS, Kelley ST, Virgin HW 4th. Murine noroviruses comprising a single genogroup exhibit biological diversity despite limited sequence divergence. J Virol. 2007 Oct;81(19):10460–73.

27. Ingle H, Makimaa H, Aggarwal S, Deng H, Foster L, Li Y, Kennedy EA, Peterson ST, Wilen CB, Lee S, Suthar MS, Baldridge MT. IFN-λ derived from nonsusceptible enterocytes acts on tuft cells to limit persistent norovirus. Sci Adv. 2023 Sep 15;9(37):eadi2562

28. Wobus CE. The Dual Tropism of Noroviruses. J Virol. 2018 Jul 31;92(16):e01010–17.

29. Passalacqua KD, Lu J, Goodfellow I, Kolawole AO, Arche JR, Maddox RJ, Carnahan KE, O’Riordan MXD, Wobus CE. Glycolysis Is an Intrinsic Factor for Optimal Replication of a Norovirus. mBio. 2019 Mar 12;10(2):e02175–18.

30. Katt WP, Cerione RA. Glutaminase regulation in cancer cells: a druggable chain of events. Drug Discov Today. 2014 Apr;19(4):450–7.

31. Wang L, Li JJ, Guo LY, Li P, Zhao Z, Zhou H, Di LJ. Molecular link between glucose and glutamine consumption in cancer cells mediated by CtBP and SIRT4. Oncogenesis. 2018 Mar 13;7(3):26.

32. Walker MC, van der Donk WA. The many roles of glutamate in metabolism. J Ind Microbiol Biotechnol. 2016 Mar;43(2-3):419–30.

33. Gualdoni GA, Mayer KA, Kapsch AM, Kreuzberg K, Puck A, Kienzl P, Oberndorfer F, Frühwirth K, Winkler S, Blaas D, Zlabinger GJ, Stöckl J. Rhinovirus induces an anabolic reprogramming in host cell metabolism essential for viral replication. Proc Natl Acad Sci U S A. 2018 Jul 24;115(30):E7158–E7165.

34. Sanchez EL, Carroll PA, Thalhofer AB, Lagunoff M. Latent KSHV Infected Endothelial Cells Are Glutamine Addicted and Require Glutaminolysis for Survival. PLoS Pathog. 2015 Jul 21;11(7):e1005052.

35. Chambers JW, Maguire TG, Alwine JC. Glutamine metabolism is essential for human cytomegalovirus infection. J Virol. 2010 Feb;84(4):1867–73.

36. Sanchez EL, Pulliam TH, Dimaio TA, Thalhofer AB, Delgado T, Lagunoff M. Glycolysis, Glutaminolysis, and Fatty Acid Synthesis Are Required for Distinct Stages of Kaposi’s Sarcoma-Associated Herpesvirus Lytic Replication. J Virol. 2017 Apr 28;91(10):e02237–16.

37. Cheng ML, Chien KY, Lai CH, Li GJ, Lin JF, Ho HY. Metabolic Reprogramming of Host Cells in Response to Enteroviral Infection. Cells. 2020 Feb 18;9(2):473.

38. Clark SA, Vazquez A, Furiya K, et al. Rewiring of the Host Cell Metabolome and Lipidome during Lytic Gammaherpesvirus Infection Is Essential for Infectious-Virus Production. Journal of Virology. 2023 Jun;97(6):e0050623.

39. Darnelle JE Jr, Eagle H. Glucose and glutamine in poliovirus production by HeLa cells. Virology. 1958 Oct;6(2):556–66.

40. Rubin H. Deprivation of glutamine in cell culture reveals its potential for treating cancer. Proc Natl Acad Sci U S A. 2019 Apr 2;116(14):6964–6968.

41. Gwangwa, M.V., Joubert, A.M. & Visagie, M.H. Effects of glutamine deprivation on oxidative stress and cell survival in breast cell lines. Biol Res 52, 15 (2019).

42. Abramowicz A, Widłak P, Pietrowska M. Different Types of Cellular Stress Affect the Proteome Composition of Small Extracellular Vesicles: A Mini Review. Proteomes. 2019 May 23;7(2):23.

43. Zhao L, Huang Y, Tian C, Taylor L, Curthoys N, Wang Y, Vernon H, Zheng J. Interferon-α regulates glutaminase 1 promoter through STAT1 phosphorylation: relevance to HIV-1 associated neurocognitive disorders. PLoS One. 2012;7(3):e32995.

44. Wang Z, Sun X, Zhao Y, Guo X, Jiang H, Li H, Gu Z. Evolution of gene regulation during transcription and translation. Genome Biol Evol. 2015 Apr 14;7(4):1155–67.

45. Katt WP, Lukey MJ, Cerione RA. A tale of two glutaminases: homologous enzymes with distinct roles in tumorigenesis. Future Med Chem. 2017 Jan;9(2):223–243.

46. Allonso D, Andrade IS, Conde JN, Coelho DR, Rocha DC, da Silva ML, Ventura GT, Silva EM, Mohana-Borges R. Dengue Virus NS1 Protein Modulates Cellular Energy Metabolism by Increasing Glyceraldehyde-3-Phosphate Dehydrogenase Activity. J Virol. 2015 Dec;89(23):11871–83.

47. Mikaeloff F, Svensson Akusjärvi S, Ikomey GM, Krishnan S, Sperk M, Gupta S, Magdaleno GDV, Escós A, Lyonga E, Okomo MC, Tagne CT, Babu H, Lorson CL, Végvári Á, Banerjea AC, Kele J, Hanna LE, Singh K, de Magalhães JP, Benfeitas R, Neogi U. Trans cohort metabolic reprogramming towards glutaminolysis in long-term successfully treated HIV-infection. Commun Biol. 2022 Jan 11;5(1):27.

48. Barrero CA, Datta PK, Sen S, Deshmane S, Amini S, Khalili K, Merali S. 2013. HIV-1 Vpr modulates macrophage metabolic pathways: a SILAC based quantitative analysis. PLoS One 8:e68376.

49. Thai, M., Thaker, S., Feng, J. et al. MYC-induced reprogramming of glutamine catabolism supports optimal virus replication. Nat Commun 6, 8873 (2015).

50. Ripoli M, D’Aprile A, Quarato G, Sarasin-Filipowicz M, Gouttenoire J, Scrima R, Cela O, Boffoli D, Heim MH, Moradpour D, Capitanio N, Piccoli C. 2010. Hepatitis C virus-linked mitochondrial dysfunction promotes hypoxia-inducible factor 1 alpha-mediated glycolytic adaptation. J Virol 84:647–660.

51. Lévy PL, Duponchel S, Eischeid H, Molle J, Michelet M, Diserens G, Vermathen M, Vermathen P, Dufour JF, Dienes HP, Steffen HM, Odenthal M, Zoulim F, Bartosch B. Hepatitis C virus infection triggers a tumor-like glutamine metabolism. Hepatology. 2017 Mar;65(3):789–803.

52. He ST, Lee DY, Tung CY, Li CY, Wang HC. Glutamine Metabolism in Both the Oxidative and Reductive Directions is Triggered in Shrimp Immune Cells (Hemocytes) at the WSSV Genome Replication Stage to Benefits Virus Replication. Front Immunol (2019) 10:2102.

53. Chen IT, Lee DY, Huang YT, Kou GH, Wang HC, Chang GD, Lo CF. 2016. Six hours after infection, the metabolic changes induced by WSSV neutralize the host’s oxidative stress defenses. Sci Rep 6:27732.

54. Keshavarz, M., Solaymani-Mohammadi, F., Namdari, H. et al. Metabolic host response and therapeutic approaches to influenza infection. Cell Mol Biol Lett 25, 15 (2020).

55. Smallwood HS, Duan S, Morfouace M, Rezinciuc S, Shulkin BL, Shelat A, Zink EE, Milasta S, Bajracharya R, Oluwaseum AJ, Roussel MF, Green DR, Pasa-Tolic L, Thomas PG. 2017. Targeting metabolic reprogramming by influenza infection for therapeutic intervention. Cell Rep 19:1640 – 1653.

56. Li Y, Webster-Cyriaque J, Tomlinson CC, Yohe M, Kenney S. Fatty acid synthase expression is induced by the Epstein-Barr virus immediate-early protein BRLF1 and is required for lytic viral gene expression. J Virol. 2004 Apr;78(8):4197–206.

57. Shin HJ, Park YH, Kim SU, Moon HB, Park DS, Han YH, Lee CH, Lee DS, Song IS, Lee DH, Kim M, Kim NS, Kim DG, Kim JM, Kim SK, Kim YN, Kim SS, Choi CS, Kim YB, Yu DY. Hepatitis B virus X protein regulates hepatic glucose homeostasis via activation of inducible nitric oxide synthase. J Biol Chem. 2011 Aug 26;286(34):29872–81.

58. Viola, A., Munari, F., Scolaro, T., & Castegna, A. (2019). The Metabolic Signature of Macrophage Responses. Frontiers in Immunology, 10, 466337.

59. Escoll P, Song OR, Viana F, Steiner B, Lagache T, Olivo-Marin JC, Impens F, Brodin P, Hilbi H, Buchrieser C. Legionella pneumophila Modulates Mitochondrial Dynamics to Trigger Metabolic Repurposing of Infected Macrophages. Cell Host Microbe. 2017 Sep 13;22(3):302–316.e7.

60. Stavru F, Bouillaud F, Sartori A, Ricquier D, Cossart P. Listeria monocytogenes transiently alters mitochondrial dynamics during infection. Proc Natl Acad Sci U S A. 2011 Mar 1;108(9):3612–7.

61. Xavier MN, Winter MG, Spees AM, den Hartigh AB, Nguyen K, Roux CM, Silva TM, Atluri VL, Kerrinnes T, Keestra AM, Monack DM, Luciw PA, Eigenheer RA, Bäumler AJ, Santos RL, Tsolis RM. PPARγ-mediated increase in glucose availability sustains chronic Brucella abortus infection in alternatively activated macrophages. Cell Host Microbe. 2013 Aug 14;14(2):159–70.

62. Bichiou, H., Bouabid, C., & Rabhi, I. (2021). Transcription Factors Interplay Orchestrates the Immune-Metabolic Response of Leishmania Infected Macrophages. Frontiers in Cellular and Infection Microbiology, 11, 660415.

63. Graziano VR, Walker FC, Kennedy EA, Wei J, Ettayebi K, Strine MS, Filler RB, Hassan E, Hsieh LL, Kim AS, Kolawole AO, Wobus CE, Lindesmith LC, Baric RS, Estes MK, Orchard RC, Baldridge MT, Wilen CB. CD300lf is the primary physiologic receptor of murine norovirus but not human norovirus. PLoS Pathog. 2020 Apr 6;16(4):e1008242.

64. Thackray LB, Wobus CE, Chachu KA, Liu B, Alegre ER, Henderson KS, Kelley ST, Virgin HWt. 2007. Murine noroviruses comprising a single genogroup exhibit biological diversity despite limited sequence divergence. J Virol 81:10460–73.

65. Gonzalez-Hernandez MB, Bragazzi Cunha J, Wobus CE. Plaque assay for murine norovirus. J Vis Exp. 2012 Aug 22;(66):e4297.

66. Chatot CL, Lawry JR, Germain B, Ziomek CA. Analysis of glutaminase activity and RNA expression in preimplantation mouse embryos. Mol Reprod Dev. 1997 Jul;47(3):248–54.

67. Allen, C.N.S.; Arjona, S.P.; Santerre, M.; Sawaya, B.E. Hallmarks of Metabolic Reprogramming and Their Role in Viral Pathogenesis. Viruses 2022, 14, 602.

68. Mullen PJ, Garcia G Jr, Purkayastha A, Matulionis N, Schmid EW, Momcilovic M, Sen C, Langerman J, Ramaiah A, Shackelford DB, Damoiseaux R, French SW, Plath K, Gomperts BN, Arumugaswami V, Christofk HR. SARS-CoV-2 infection rewires host cell metabolism and is potentially susceptible to mTORC1 inhibition. Nat Commun. 2021 Mar 25;12(1):1876.

69. Mullen AR, Wheaton WW, Jin ES, Chen PH, Sullivan LB, Cheng T, Yang Y, Linehan WM, Chandel NS, DeBerardinis RJ. Reductive carboxylation supports growth in tumour cells with defective mitochondria. Nature. 2011 Nov 20;481(7381):385–8.

70. Yang L, Achreja A, Yeung TL, Mangala LS, Jiang D, Han C, Baddour J, Marini JC, Ni J, Nakahara R, Wahlig S, Chiba L, Kim SH, Morse J, Pradeep S, Nagaraja AS, Haemmerle M, Kyunghee N, Derichsweiler M, Plackemeier T, Mercado-Uribe I, Lopez-Berestein G, Moss T, Ram PT, Liu J, Lu X, Mok SC, Sood AK, Nagrath D. Targeting Stromal Glutamine Synthetase in Tumors Disrupts Tumor Microenvironment-Regulated Cancer Cell Growth. Cell Metab. 2016 Nov 8;24(5):685–700.

71. Zhao H, Yang L, Baddour J, Achreja A, Bernard V, Moss T, Marini JC, Tudawe T, Seviour EG, San Lucas FA, Alvarez H, Gupta S, Maiti SN, Cooper L, Peehl D, Ram PT, Maitra A, Nagrath D. Tumor microenvironment derived exosomes pleiotropically modulate cancer cell metabolism. Elife. 2016 Feb 27;5:e10250.

72. Zhu Z, Achreja A, Meurs N, Animasahun O, Owen S, Mittal A, Parikh P, Lo TW, Franco-Barraza J, Shi J, Gunchick V, Sherman MH, Cukierman E, Pickering AM, Maitra A, Sahai V, Morgan MA, Nagrath S, Lawrence TS, Nagrath D. Tumour-reprogrammed stromal BCAT1 fuels branched-chain ketoacid dependency in stromal-rich PDAC tumours. Nat Metab. 2020 Aug;2(8):775–792.

73. Achreja A, Yu T, Mittal A, Choppara S, Animasahun O, Nenwani M, Wuchu F, Meurs N, Mohan A, Jeon JH, Sarangi I, Jayaraman A, Owen S, Kulkarni R, Cusato M, Weinberg F, Kweon HK, Subramanian C, Wicha MS, Merajver SD, Nagrath S, Cho KR, DiFeo A, Lu X, Nagrath D. Metabolic collateral lethal target identification reveals MTHFD2 paralogue dependency in ovarian cancer. Nat Metab. 2022 Sep;4(9):1119–1137.

74. Strelko CL, Lu W, Dufort FJ, Seyfried TN, Chiles TC, Rabinowitz JD, Roberts MF. Itaconic acid is a mammalian metabolite induced during macrophage activation. J Am Chem Soc. 2011 Oct 19;133(41):16386–9.

75. Michelucci A, Cordes T, Ghelfi J, Pailot A, Reiling N, Goldmann O, Binz T, Wegner A, Tallam A, Rausell A, Buttini M, Linster CL, Medina E, Balling R, Hiller K. Immune-responsive gene 1 protein links metabolism to immunity by catalyzing itaconic acid production. Proc Natl Acad Sci U S A. 2013 May 7;110(19):7820–5.

76. O’Neill LAJ, Artyomov MN. Itaconate: the poster child of metabolic reprogramming in macrophage function. Nat Rev Immunol. 2019 May;19(5):273–281.

77. Cordes T, Wallace M, Michelucci A, Divakaruni AS, Sapcariu SC, Sousa C, Koseki H, Cabrales P, Murphy AN, Hiller K, Metallo CM. Immunoresponsive Gene 1 and Itaconate Inhibit Succinate Dehydrogenase to Modulate Intracellular Succinate Levels. J Biol Chem. 2016 Jul 1;291(27):14274–14284.

78. Wobus CE, Peiper AM, McSweeney AM, Young VL, Chaika M, Lane MS, Lingemann M, Deerain JM, Strine MS, Alfajaro MM, Helm EW, Karst SM, Mackenzie JM, Taube S, Ward VK, Wilen CB. Murine Norovirus: Additional Protocols for Basic and Antiviral Studies. Curr Protoc. 2023 Jul;3(7):e828.

79. Smith BJ. SDS Polyacrylamide Gel Electrophoresis of Proteins. Methods Mol Biol. 1984;1:41–55.

80. Taciak B, Białasek M, Braniewska A, Sas Z, Sawicka P, Kiraga Ł, Rygiel T, Król M. Evaluation of phenotypic and functional stability of RAW 264.7 cell line through serial passages. PLoS One. 2018 Jun 11;13(6):e0198943.

81. Nabeel Attarwala, Cissy Zhang, Anne Lee. Diseases & Disorders | Therapies Targeting Glutamine Addiction in Cancer. Joseph Jez, editor. Encyclopedia of Biological Chemistry III (Third Edition), Elsevier; 2021. pp. 452–461.

82. Sosnovtsev SV, Belliot G, Chang KO, Prikhodko VG, Thackray LB, Wobus CE, Karst SM, Virgin HW, Green KY. Cleavage map and proteolytic processing of the murine norovirus nonstructural polyprotein in infected cells. J Virol. 2006 Aug;80(16):7816–31.

83. Hyde JL, Mackenzie JM. Subcellular localization of the MNV-1 ORF1 proteins and their potential roles in the formation of the MNV-1 replication complex. Virology. 2010 Oct 10;406(1):138–48.

84. Jahun AS, Sorgeloos F, Chaudhry Y, Arthur SE, Hosmillo M, Georgana I, Izuagbe R, Goodfellow IG. Leaked genomic and mitochondrial DNA contribute to the host response to noroviruses in a STING-dependent manner. Cell Rep. 2023 Mar 28;42(3):112179.

85. Baker E. Characterization of the NS1-2 and NS4 proteins of murine norovirus: PhD Thesis. University of Otago, Microbiology & Immunology; 2012.

